# Biomimetic fractal topography enhances podocyte maturation in vitro

**DOI:** 10.1101/2024.03.16.585039

**Authors:** Chuan Liu, Praful Aggarwal, Karl T. Wagner, Shira S. Landau, Teng Cui, Xin Song, Laleh Shamaei, Naimeh Rafatian, Yimu Zhao, Sonia Rodriguez-Ramirez, Keith Morton, Elizabeth Virlee, Chen Yu Li, Dawn Bannerman, Simon Pascual-Gil, Sargol Okhovatian, Anastasia Radisic, Sergi Clotet-Freixas, Teodor Veres, Mohtada Sadrzadeh, Tobin Filleter, Ulrich Broeckel, Ana Konvalinka, Milica Radisic

## Abstract

Cells and tissues in their native environment are organized into intricate fractal structures, which are rarely recapitulated in their culture *in vitro*. The extent to which fractal patterns that resemble complex topography in vivo influence cell maturation, and the cellular responses to such shape stimulation remain inadequately elucidated. Yet, the application of fractal cues (topographical stimulation via self-similar patterns) as an external input may offer a much-needed solution to the challenge of improving the differentiated cell phenotype *in vitro*. Here, we established fractality in podocytes, branching highly differentiated kidney cells, and glomerulus structure. Biomimetic fractal patterns derived from glomerular histology were used to generate topographical (2.5-D) substrates for cell culture. Podocytes grown on fractal topography were found to express higher levels of functional markers and exhibit enhanced cell polarity. To track morphological complexities of differentiated podocytes, we employed a fluorescent labelling assay where labelled individual cells are tracked within otherwise optically silent confluent cell monolayer to reveal cell-cell interdigitation. RNAseq analysis suggests enhanced ECM deposition and remodeling in podocytes grown on fractal topography compared to flat surface or non-fractal microcurvature, mediated by YAP signaling. The incorporation of fractal topography into standard tissue culture well plates as demonstrated here may serve as a user-friendly bioengineered platform for high-fidelity cell culture.

## 1. Introduction

Podocytes are highly specialized, terminally differentiated epithelial cells in the kidney glomerulus that cover the outer surfaces of glomerular capillaries and form a selective barrier for blood filtration. Podocytes possess multi-scale foot processes branching from the main cell body, which compose slit diaphragms that serve as a filtration barrier to prevent macromolecules such as proteins from passing through, ensuring that macromolecules are retained in the bloodstream. During development, podocytes’ morphology gradually becomes complex as their foot processes turn from a primitive to a mature state.^1^ This intricate, branching morphology is an important indicator of podocyte health and function as the deterioration or diminishment of this interdigitating pattern, referred to as foot process effacement, leads to glomerular disease characterized by significant proteinuria.

Given the definition of fractals as geometry with recurring self-similar patterns at continuously smaller scales, the branching of foot processes at multiple levels/scales suggests that podocytes are fractal objects. As such, their morphology can be characterized by fractal dimension (Df), a ratio of the change in detail to the change in scale and often used for quantifying complex geometry. Similar to podocytes, a kidney glomerulus has a complex architecture, with fractal analysis performed on itself^2^ or its components^3^ under healthy and pathological/injury conditions. Not surprisingly, state of tissue health has been reported to correlate with their fractality,^4^ which suggests engineering microenvironmental factors such as biophysical shape cues could control cell or tissue fate.

It continues to pose a difficulty recreating physiologically pertinent podocytes, especially with human cells in culture.^5^ In addition to gene expression diverging considerably from the native conditions and phenotypes exhibiting immaturity, there is a lack of functional readouts that allow for injury modelling. Various methods have been developed to help mature cultured podocytes or generate more advanced glomerulus models *in vitro*, including biochemical cues^6^, organoid differentiation^7^, decoration of substrates with extracellular matrix (ECM)^8^, cyclic mechanical stimulation^9^, and flow with vascularization^9–11^. However, mimicking the complex glomerular architecture has not yet been accomplished. Flat surfaces or membranes are utilized for cell and tissue culture, even in microfluidic systems. On the other hand, it is well known that topographical cues play an important role in cell behaviour. Specifically, simple round-shaped topography or microcurvature has been shown to improve the fidelity of podocytes cultured *in vitro*.^11,12^ Building upon the idea of shape stimulation^13–15^, we were interested in investigating whether more biomimetic patterns could further enhance podocyte maturation. Here, we further delineated fractality of podocyte and glomerulus structure and explored the use of glomerulus-mimicking fractal topography to support higher fidelity podocyte culture *in vitro*.

## 2. Results

### 2.1. Native glomerulus and podocyte demonstrate fractal patterns

A typical fractal shape shows self-similarity over multiple scales of measurement, such as the famous Koch snowflake, whose size expands 4 times as the scale decreases by 1/3, corresponding to a fractal dimension of 1.26 in theory (**Fig. 1a**). Both the glomerulus and podocyte demonstrate a complex structure under X-ray nanotomography and focused ion beam/scanning electron microscopy (SEM), respectively, suggesting fractal dimension as a suitable index for the quantification of their geometry (**Fig. 1b**). Preliminary fractal analysis of histology images of glomeruli from healthy and pathological samples also showed the existence of fractality and a change in fractal dimension between healthy and diseased states (**Fig. 1c**).

**Fig. 1.**
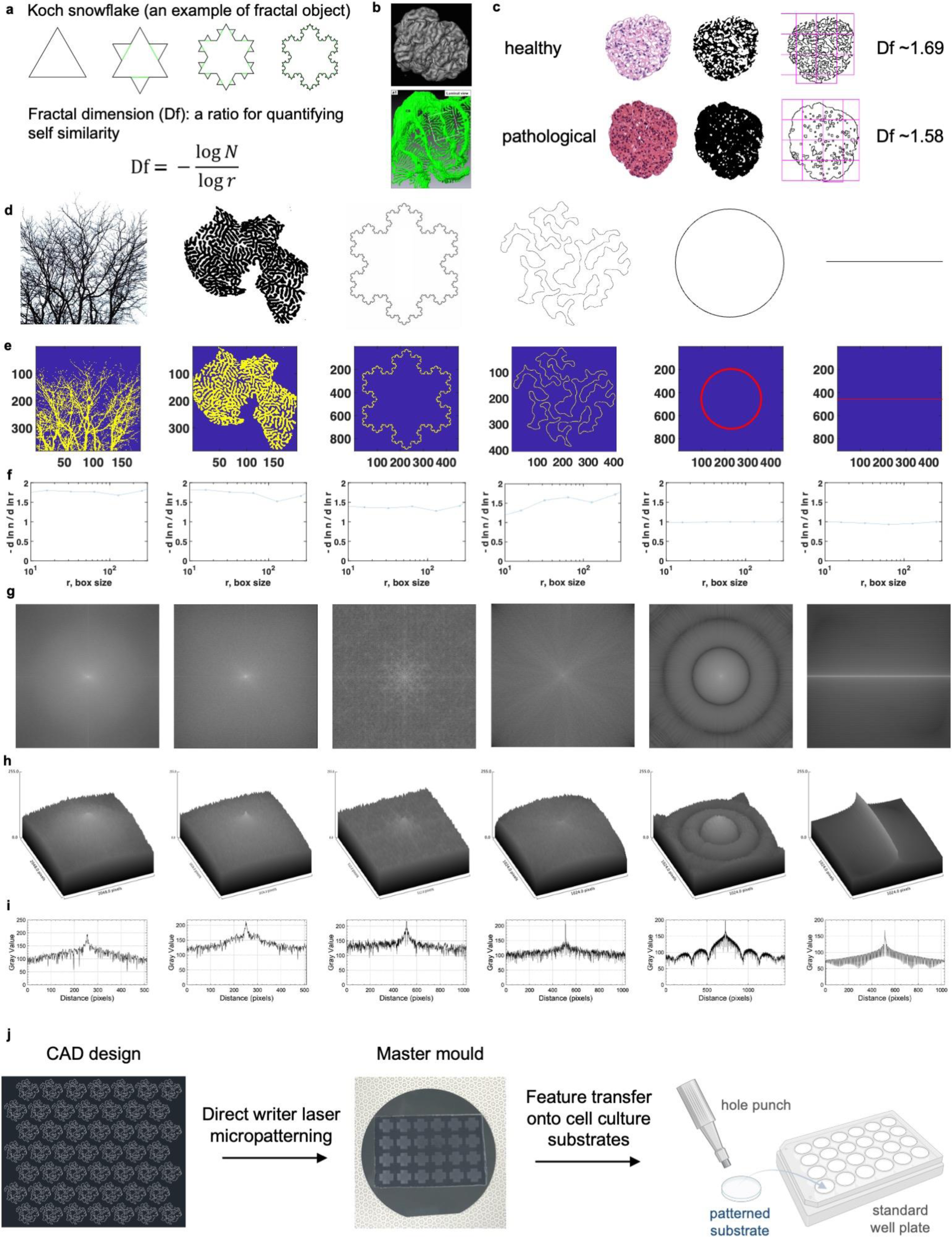
Fractal metrics characterize architecture of podocytes and glomeruli in healthy and diseased states. **a,** Koch snowflake is a typical example of fractal shape, which can be characterized by fractal dimension (Df). **b,** Cast of a glomerulus from X-ray nanotomography, reproduced with permission,^17^ and a single podocyte reconstructed from serial focused ion beam/scanning electron microscope (FIB/SEM) images, reproduced with permission.^18^ **c,** Original images, thresholded masks, and projected traces of cross-sectional H&E histology, taken from healthy and pathological (proliferative lesions) sections. Histology reproduced with permission.^19^ **d-i,** Characterization of fractal properties in a podocyte^16^ and a glomerulus in comparison to standard fractal (a tree, a Koch snowflake) and non-fractal (a line and a circle) controls. **d,** Original images of the different shapes (from left to right): tree, podocyte (reproduced with permission^16^), Koch snowflake, glomerular trace^13–15^, circle, line. **e,** Binary images of different shapes with the background removed (unit: pixel). Original thickness of a circle and a line is 2 pixels, thick red color added for visualization. **f,** The fractal dimension range graphed over the box size, calculated by the box-counting method. **g,** FFT of the various shapes. **h,** Surface plotting of the FFT results. **i,** A profile plot through the center of the FFT surface. HF, high fractal. FFT, fast Fourier transform. **j,** Generation of topographical substrates with glomerulus-mimicking pattern from design to fabrication.

Previously, histological glomerular tracing was developed and reported by Korolj et al.^13–15^ to obtain the outline of a glomerulus from histology images. To confirm the occurrence of fractality, we compared the planar projection of an SEM image of a podocyte^16^ (**Fig. 1d**) and a glomerular trace^13–15^ (**Fig. 1d**) to objects that are widely accepted to be fractal, specifically a branching tree and Koch snowflake, a common fractal computational control (**Fig. 1d**). We contrasted these to the objects that are known not to be fractal, specifically the perfect circle and the straight line (**Fig. 1d**). Relying on the box counting method we determined that both the mature podocyte and the outline of a glomerulus exhibit multi-fractality, similar to that of a branching tree and of the Koch snowflake (**Fig. 1f**). Whereas the non-fractal objects, a circle and a line, exhibited a Df=1 regardless of the box size, the podocyte (Df ranging from 1.52 to 1.82) and the glomerulus (Df ranging from 1.31 to 1.73) exhibited a Df similar to values observed for the tree (Df from 1.68 to 1.80) and the Koch snowflake (Df from 1.27 to 1.39).

Importantly, whereas dominant frequencies were easily identifiable for the circle and the line (**Fig. 1g-i**) upon fast Fourier transformation (FFT), no dominant frequency was apparent for the podocyte, the glomerulus, a tree and the Koch snowflake (**Fig. 1g-i**). Together with the fractal analysis (**Fig. 1c-i**), these results further confirm the presence of fractality both in the adult, mature, podocyte shape, as well as histological outline of the glomerulus.

In the remainder of the manuscript, we focus on a single Df value, to balance experimental feasibility of constructing physical scaffolds of clearly delineated patterning categories with the biology of the cells cultured thereon.

### 2.2. Fractal topography increases expression of slit diaphragm markers in mouse podocytes

To investigate the effect of fractal topography on podocyte culture *in vitro*, we used topographical cell culture substrates with biomimetic fractal patterns derived from the podocyte microenvironment observed in histology slices from healthy and pathological samples, via an approach previously reported by Korolj et al.^13–15^ The design unit of the biomimetic fractal pattern from a healthy histology image was confirmed to be fractal by box counting fractal analysis (**Fig. 1f**) and packed across a 2D surface to form a patterned area of desirable size. These fractal patterns were then transferred to a master mold via a laser direct writer, which was used to fabricate fractal topographical substrates made of polydimethylsiloxane (PDMS). To incorporate the patterned substrates into the standard 2D cell culture workflow, we punched them into circular inserts of the size of a 24-well and inserted them into a standard tissue culture well plate (**Fig. 1j**).

The biomimetic fractal topography corresponding to a healthy state created by histological glomerular tracing^13–15^ was referred to as high-fractal (HF) topography and each HF design unit was considered a glomerulus slice with a diameter of approximately 150 µm. As the biomimetic fractal topography derived from a pathological histology image using the same method^13–15^ had a lower Df, it was referred to as low-fractal (LF) topography. To dissect the effect of fractality on podocytes, a round topography (RT) made of an array of microhemispheres (each with the diameter of a glomerulus slice), previously developed and reported,^13–15^ was used as a non-fractal control, in addition to the simple flat substrates with no topography (NT).^15^ The surface features of the substrates from these different groups can be clearly visualized under SEM and 3D digital microscopy (**Fig. 2a**), with an increasing Df from the non-fractal RT control group to the fractal LF and HF groups as expected (**Fig. 2c**).

**Fig. 2.**
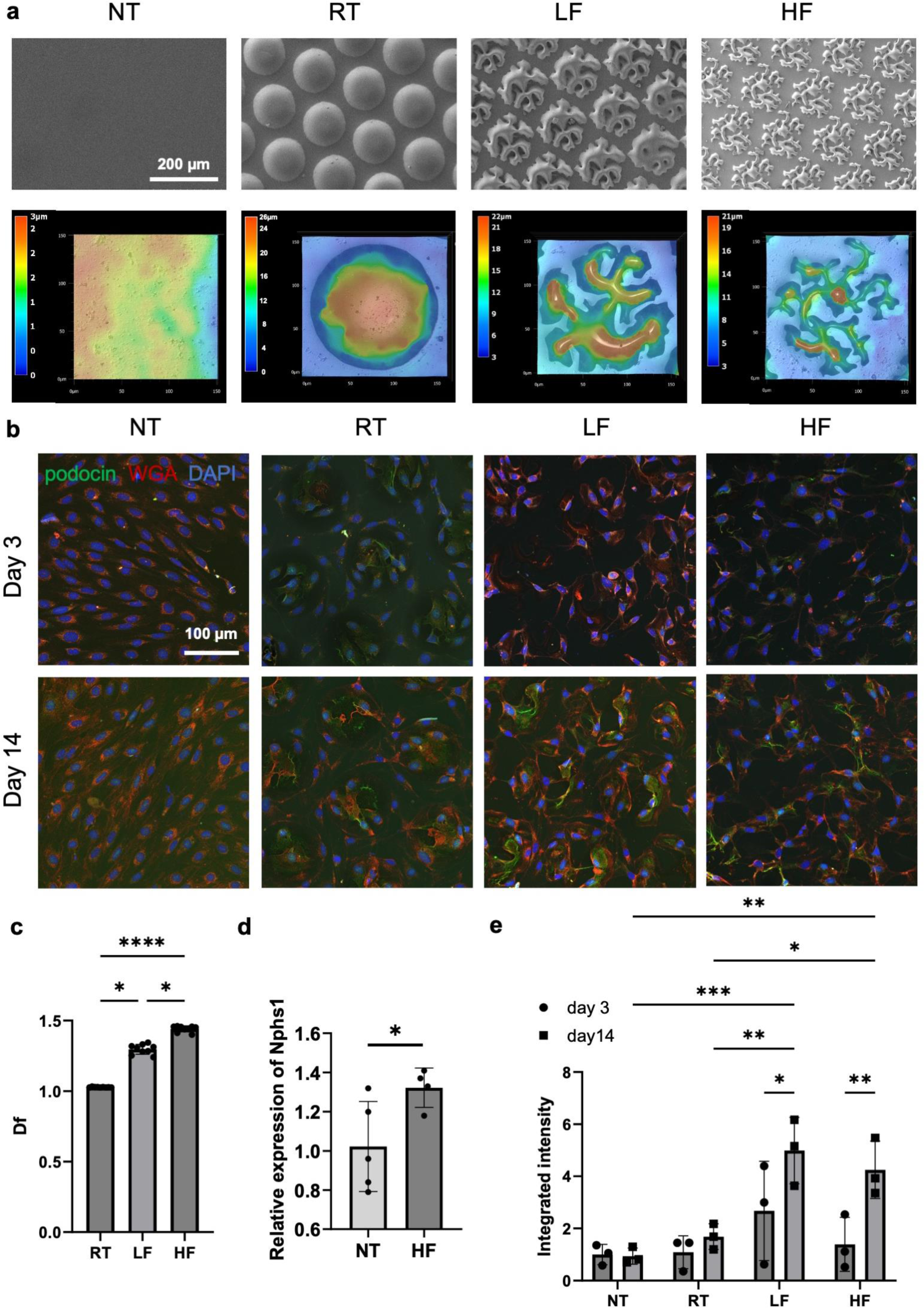
Fractal topography increases expression of slit diaphragm markers in mouse podocytes. **a,** Scanning electron micrographs (SEM) and 3D display of digital micrographs of PDMS scaffolds showing successful transfer of non-topographic (NT), round non-fractal (RT), and biomimetic low and high fractal (LF and HF) patterns to substrates. **b-c,** Expression of slit diaphragm protein podocin increases on fractal substrates with time in culture. **b,** Immunostaining of podocin (counterstained with WGA) from mouse podocytes grown on NT, RT, LF and HF substrates and differentiated for 3 and 14 days. **c,** Fractal dimension of RT, LF, and HF patterns from 3 topographical regions on n=3 scaffold samples. One-way ANOVA with Tukey’s multiple comparisons test performed. Data plotted as mean ± SD. **d,** Relative gene expression of *Nphs1* to *Gapdh* in mouse podocytes cultivated on n=5 NT and n=4 HF scaffolds. Data normalized to NT. Student’s t-test for statistical testing. Data plotted as mean ± SD. **e,** Quantification of podocin immunofluorescence signal. n=3 scaffold samples. Data normalized to day 3 NT. Two-way ANOVA with Tukey’s multiple comparisons test performed. Data plotted as mean ± SD. *p≤0.05, **p≤0.01, ***p≤0.001, ****p≤0.0001. NT, no topography; RT, round topography; LF, low fractal; HF, high fractal.

A conditionally immortalized mouse podocyte cell line, E11, is commonly used for studying podocyte biology since human podocyte cells are scarce. The conditional immortalization preserves podocytes’ terminal differentiation state while allowing for cell expansion. We assessed the gene expression of a slit diaphragm marker, *Nphs1*, from E11 podocytes grown on NT and HF substrates by quantitative polymerase chain reaction (qPCR) and found a significantly enhanced expression on HF compared to the NT control (**Fig. 2d**).

Next, we evaluated the expression of another slit diaphragm marker, podocin, from podocytes cultured on flat (NT) and topographical (RT, LF, HF) substrates over time. E11 podocytes have a thermoswitching mechanism that could deactivate the forced cell cycle processes and switch from a proliferation to differentiation mode. A cultivation period of at least 14 days is commonly used and regarded as required for sufficient differentiation. As such, we selected day 3 and day 14 as the early (immature) and late (mature) endpoints, respectively. For true interdigitation, cell-cell contact is needed. Thus, structures that we are focusing on can only be reliably observed once the cells are approaching confluence and beyond. Cells cultivated on NT substrates tend to proliferate more, thus achieving conditions for formation of inter-digitations faster (**Fig. 2b**). Although in early cultures (Day 3), HF substrates did not appear to have substantial podocin presence (**Fig. 2b-c**), as the culture time increased fractal substrates exhibited a clear improvement of podocin presence over flat substrates, whereas cells on flat substrates exhibited levels as low as on Day 3 even when cell culture was taken to Day 14 (**Fig. 2b-c**).

The biomimetic fractal topography not only promoted podocin expression, but also guided cytoskeleton arrangement (**Supplementary Fig. 1**) when E11 podocytes were seeded on substrates with both flat surfaces and fractal patterns, and differentiated for 14 days. There was a visible difference in f-actin alignment between cells grown on flat surfaces and those grown on fractal topography where f-actin was elongated and aligned on flat surfaces while exhibiting a more branched, in vivo-like pattern on fractal topography.

### 2.3. Fractal topography elicits enhanced signs of cell polarity in mouse podocytes

Apical surfaces of epithelial cells are decorated with glycans that contribute to regulation of cell structure, function, and communication via protruding forms such as charged villi, blebs, and secreted extracellular vesicles (EVs).^20^ We hypothesized that the arrangement of ECM proteins, facilitated by fractal topographical cues, could mediate assembly of podocyte structures involved in polarization. SEM underscored the interdigitated 3D morphology of cells on fractal substrates vs flat and RT controls (**Fig. 3a**). The protruding apical structures appeared to progress from rounded (mushroom regime) in the NT group to elongated finger-like protrusions (brush regime) in the HF group, according to the glycocalyx development model by Shurer *et al*. (**Fig. 3a**).^20^

**Fig. 3.**
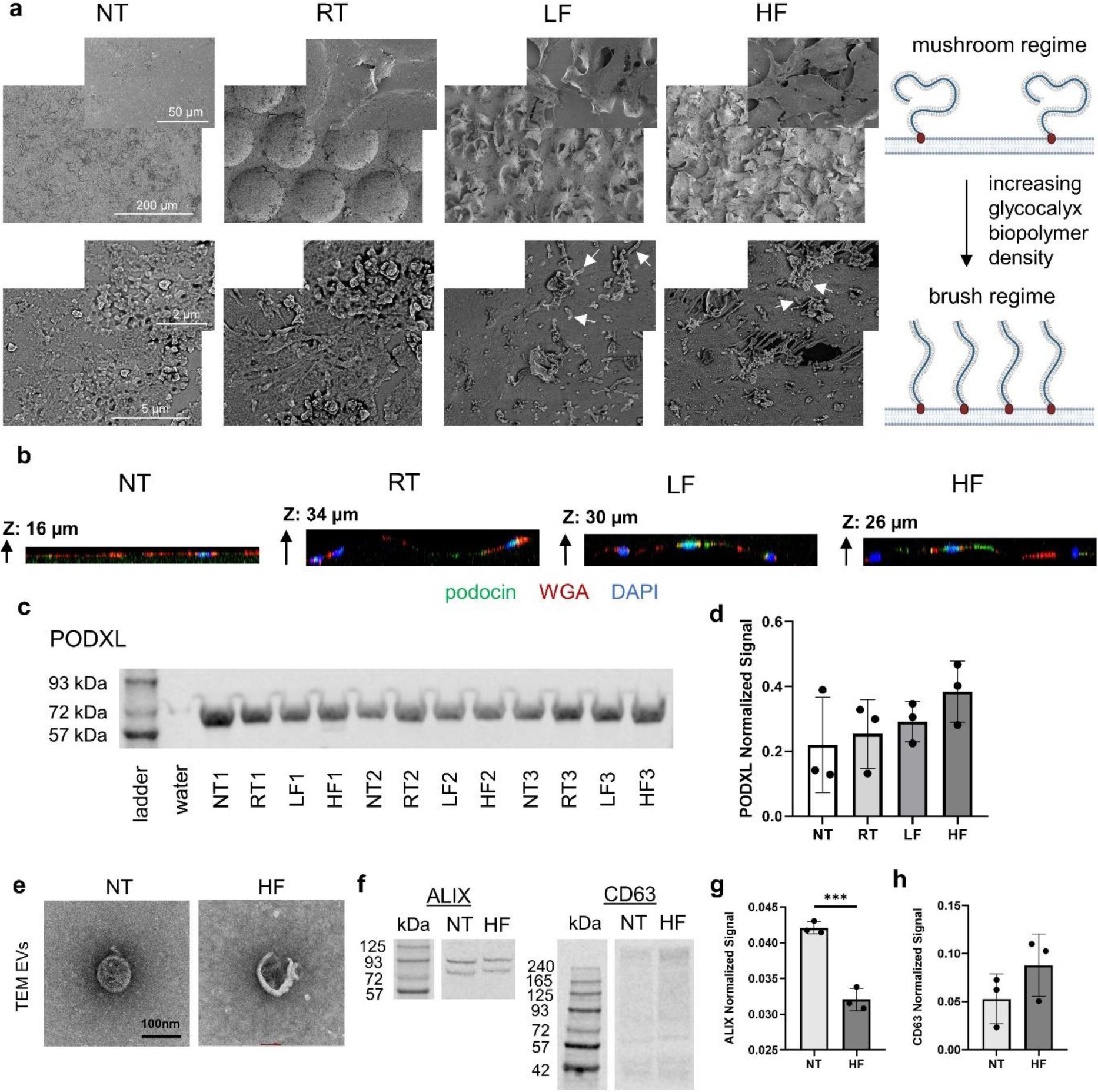
Fractal topographic patterning promotes signs of polarity in mouse podocytes. **a,** SEM of podocytes cultivated on NT, RT, LF and HF scaffolds from low to high magnifications. Arrows point to finger-like apical protrusions. **b,** Immunofluorescence images of podocytes cultivated on NT, RT, LF and HF scaffolds showing podocin (green) presence in z-stacks. **c,** Western blot of podocalyxin from podocytes cultivated on NT, RT, LF and HF scaffolds. **d,** Quantification of PODXL signal from western blot (normalized to GAPDH). n=3 scaffold samples. One-way ANOVA statistical testing shows no significant difference. **e,** TEM of EVs isolated from culture media of podocytes cultivated on NT and HF scaffolds. **f,** Western blot of ALIX and CD63 from EV samples isolated from culture media of mouse podocytes cultivated on NT and HF scaffolds. **g-h,** Quantification of **g,** ALIX and **h,** CD63 signal from western blot (normalized to total protein). n=3 scaffold samples. Student t-test used for statistical testing. Data plotted as mean ± SD. ***p≤0.001. SEM, scanning electron micrograph; TEM, transmission electron micrograph; EV, extracellular vesicle; NT, no topography; RT, round topography; LF, low fractal; HF, high fractal.

Fractal topography was critical for podocyte marker localization, as confocal z-stacks demonstrated greater portions of podocin localization on top of the topography in the fractal groups (**Fig. 3b**) as well as greater presence of podocin in the fractal groups (**Supplementary Fig. 2**). Podocalyxin, a negatively charged sialylated glycoprotein involved in polarization^21^ and basal organization,^22^ exhibited stable protein expression and an increasing trend in the HF group **(Fig. 3c-d, Supplementary Fig. 3a)**.

Epithelial polarity is often associated with a dense glycocalyx,^22,23^ and a dense glycocalyx is likewise associated with the budding of extracellular vesicles (EV).^20^ TEM revealed the cup-like shape of EV’s **(Fig. 3e)** and expression of EV markers ALIX and CD63 **(Fig. 3f-h**, full blots available in **Supplementary Fig. 3b-c)**. Although both are exosome markers, ALIX exhibited decreased expression on HF substrates while high ALIX has been reported in exosomes of certain cancers.^24,25^ Collectively, these observations point to the signs of physiological relevance of podocyte apical surface morphology on fractal substrates.^20,26^

### 2.4. Biomimetic fractal topography regulates cytoskeleton arrangement in mouse podocytes

As it is known that the wavelength of a substrate impacts biological responses^27,28^ and the sharp microcurvature of the multiscale-curved HF pattern on the lower end is not represented by the hemi-spherical non-fractal RT control, we designed a non-fractal ordered “spiky” topography (ST) surface that aimed to capture the pointy features at the tips of the HF pattern, with spikes placed at a regular distance in a lattice (**Fig. 4a-b**). To ensure all the conditions were kept the same, we fabricated the ST and HF master molds on the same wafer. The resulting ST and HF substrates have similar feature heights (**Fig. 4c-e**), with the peaks in the radius of curvature distribution overlapping at approximately 2μm (**Fig. 4f**).

**Fig. 4.**
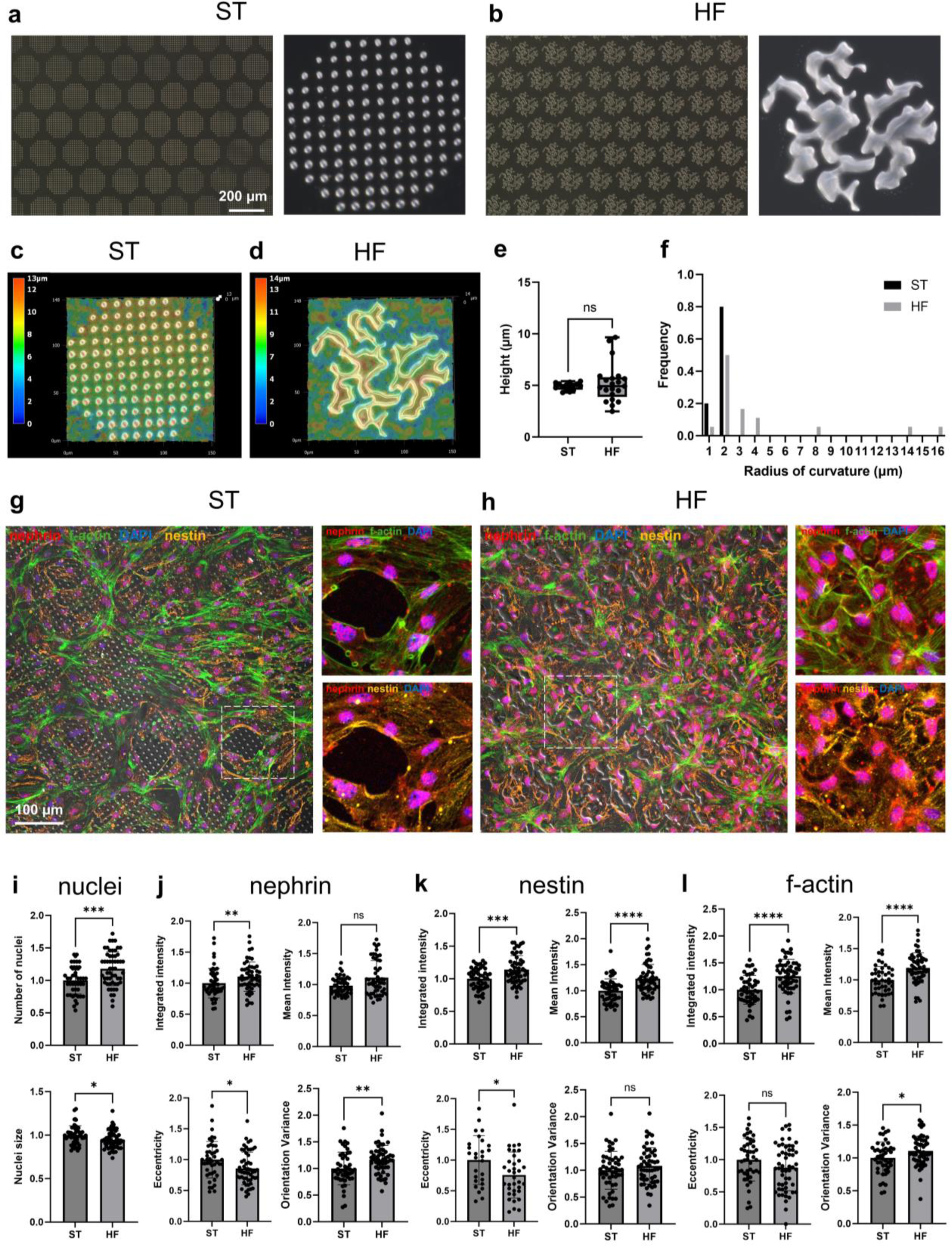
Mouse podocytes exhibit more in vivo-like morphology on biomimetic fractal topography with multiscale curvature versus equally sharp non-fractal ordered topography with a narrow curvature range. **a-b,** Low (left) and high (right) magnification digital microscope images of **a,** ST and **b,** HF scaffolds. **c-d,** Surface characterization of **c,** ST and **d,** HF scaffolds. **e-f,** Feature **e,** height and **f,** radius of curvature measured from regions on 3 ST and 3 HF scaffolds. **g-h,** Immunofluorescence images of mouse podocytes cultured on **g,** ST and **h,** HF scaffolds and stained for nephrin (red), nestin (orange), f-actin (green) and DAPI (blue). **i-l,** Quantification and directionality analysis of immunofluorescence signals from **i,** DAPI (nuclei number and size), **j,** nephrin (intensity and directionality), **k,** nestin (intensity and directionality), and **l,** f-actin (intensity and directionality) within the topographical regions of 4 ST and 4 HF scaffolds. Student’s t-test or Mann-Whitney nonparametric test was performed for statistical analysis. *p<0.05, **p<0.01, ***p<0.001, ****p<0.0001. ST, spiky topography; HF, high fractal.

Podocytes cultivated on both the ST and HF substrates formed a confluent layer; however, areas without cells (“holes”) were observed on the ST substrates (**Fig. 4g**) but not on the HF substrates (**Fig. 4h**). Thus, HF substrates support formation of a cellular layer in full confluency as reflected by the higher nuclei number in the topographical regions covering the patterns, despite a smaller nuclei size (**Fig. 4i**). Notably, we found stronger fluorescence signals of a slit diaphragm marker (nephrin) and two filament components (f-actin, nestin) in the topographical regions of HF versus ST patterns (**Fig. 4j-l**), suggesting higher expression of nephrin, f-actin, and nestin in cells cultured on the HF substrates. As an intermediate filament, nestin maintains podocyte foot process^29^ as well as actin structures,^30^ which is consistent with its increased expression in the HF group accompanied by the increased expression of nephrin and f-actin. As nestin is shown to be upregulated to protect podocytes in response to injury,^30,31^ the elevated expression of nestin on HF substrates indicates podocytes’ enhanced resilience to injury when cultured on HF topography. Interestingly, nephrin on the HF topography exhibits a lower extent of both elongation and alignment compared to that on the ST topography (**Fig. 4j**); similar results were observed for nestin with respect to alignment (**Fig. 4k**) and f-actin with respect to alignment (**Fig. 4l**). Decreased elongation and alignment of the cytoskeleton structures indicate a more in vivo-like arrangement since native podocytes possess an arborized as opposed to elongated morphology.

### 2.5. Mosaic assay reveals single cell morphology of human podocytes cultured *in vitro*

With response to fractal topographical stimulation observed in mouse podocytes, we switched to human podocyte culture for a more physiologically relevant model. Although human kidney ECM supported podocyte attachment, more reproducible results in terms of obtaining a confluent coverage with cells are obtained with a Matrigel coating, likely due to the elevated batch-to-batch variability of human ECM. (**Supplementary Fig. 4**).

It has been a challenge to observe podocyte morphology *in vitro* due to the difficulty in delineating the border of a single cell from those of its neighbors. To visualize single cell morphology in a confluent layer of podocytes cultured in vitro, optically silent podocytes were mixed with their counterpart labelled with green fluorescent protein (GFP) to obtain a mixed cell population with approximately 1/12 fluorescent cells. This mixed cell population is referred to as Mosaic assay as described previously^13–15^, enabling single cell morphology to be easily visualized under fluorescence (**Fig. 5a,d,h**). Using Mosaic assay, fractal dimension of podocyte morphology, an indicator of cell maturation and function, can be analyzed via the box counting method.

**Fig. 5.**
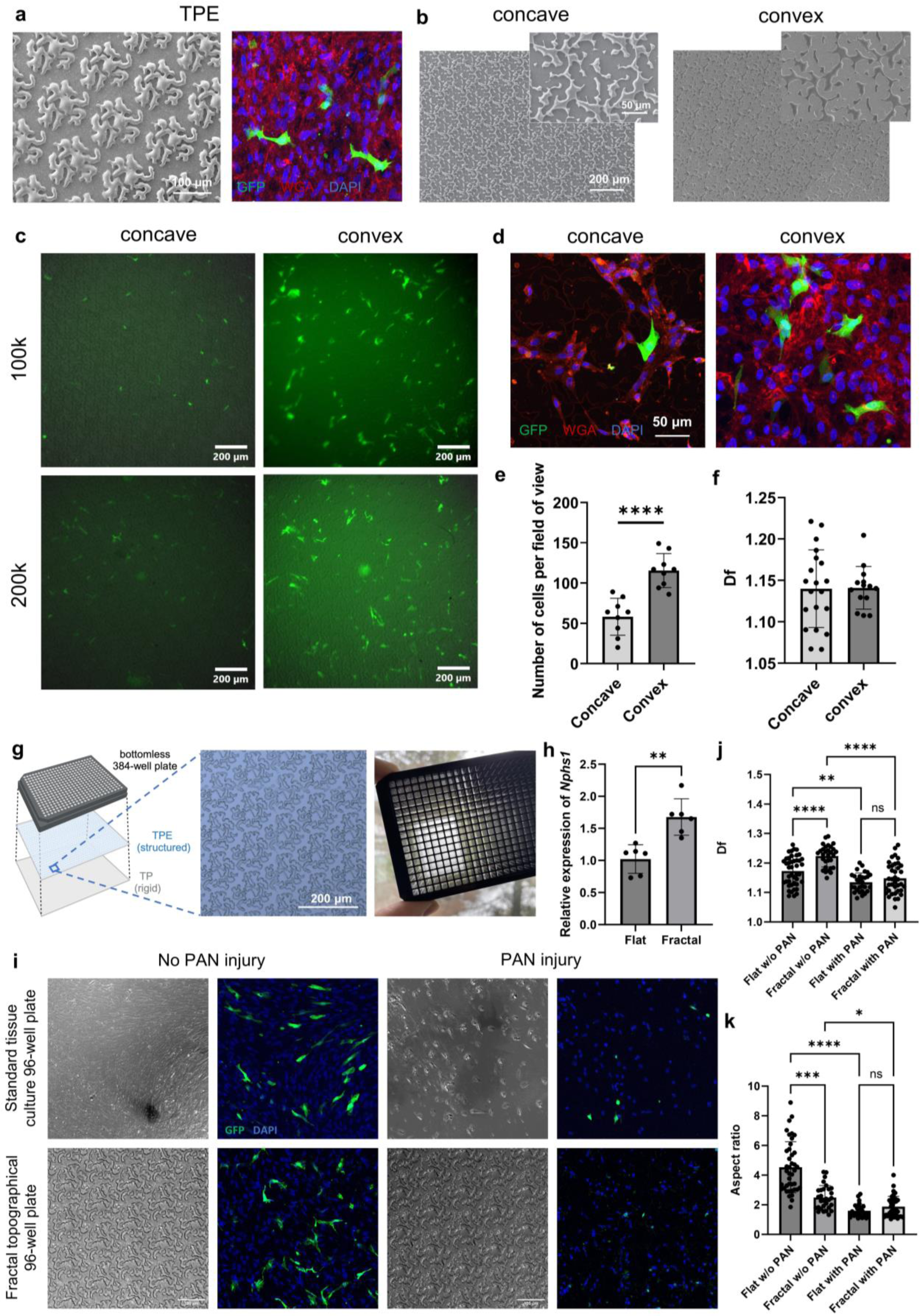
Mosaic assay reveals single cell morphology of human podocytes cultured in vitro. **a,** SEM of TPE scaffolds with HF pattern and Mosaic assay of non-conditionally immortalized human podocytes cultivated on TPE scaffolds. **b,** SEM of high density fractal PDMS substrates in concave and convex directions. Low and high magnification shown. **c-d,** Mosaic assay of non-conditionally immortalized human podocytes cultivated on concave and convex HF scaffolds (n=3) for a period of 5 days and imaged by **c,** epifluorescence (seeding densities at 100,000 and 200,000 cells/cm^2^) and **d,** confocal microscopy (seeding density at 100,000 cells/cm^2^). **e-f,** Quantification of **e,** cell numbers and **f,** whole cell morphology by fractal dimension of Mosaic assay human podocytes cultivated on concave and convex HF scaffolds (n=3) for 5 days. Student t-test used for statistical testing. Data plotted as mean ± SD. ****p≤0.0001. **g-k,** Fractal topographical well plates support higher fidelity podocyte culture compared to standard tissue culture well plates. **g,** Schematic and images of high density HF patterns covering the entire well in a 384-well format. **h,** Relative gene expression of *Nphs1* to *Gapdh* in mouse podocytes seeded and differentiated on the standard and fractal well plates. n=6 samples per group. Data normalized to flat (standard 96-well plate). Student’s t-test for statistical testing. Data plotted as mean ± SD. **i,** Brightfield and fluorescence images of Mosaic human podocytes cultured on the standard and fractal well plates, with and without PAN-induced injury. **j-k,** Quantification of cell morphology **j,** Df and **k,** aspect ratio of Mosaic human podocytes cultivated on the standard and fractal well plates, with and without PAN-induced injury. n=3 samples per group. One-way ANOVA with Tukey’s multiple comparisons test performed for **j**. Kruskal-Wallis with Dunn’s multiple comparisons test performed for **k**. Data plotted as mean ± SD. *p≤0.05, **p≤0.01, ***p≤0.001, ****p≤0.0001. GFP, green fluorescent protein. PAN, puromycin aminonucleoside. TPE, thermoplastic elastomer; HF, high fractal.

To eliminate the use of PDMS and to verify the applicability of fractal topography on alternative substrate materials with the potential of scalable production, it was necessary to demonstrate podocyte branching morphology on substrates prepared via hot embossing of a thermoplastic elastomer (TPE) and tissue culture polystyrene. The fractal features were clearly imprinted **(Fig. 5a, Supplementary Fig. 5a)** and successfully assembled with a bottomless well plate for cell culture **(Fig. 5g, Supplementary Fig. 5b)**. Human podocytes cultured on the TPE and polystyrene substrates were able to form a confluent layer, exhibiting branching morphology (**Fig. 5a, Supplementary Fig. 5c**). The use of thermoplastic materials enabled us to rapidly fabricate high density fractal patterns covering the entire well in a 24-, 96-, and 384-well format **(Fig. 5g, Supplementary Figs. 5 and 6)**.

Furthermore, we attempted to minimize the space in between slices of HF topography as it was those biophysical cues that drove the cellular response such as increased expression of podocyte markers and polarity. The HF patterns for the master mold were modified by more densely packing the HF design unit in order to reduce the flat region in between the HF design units (**Supplementary Fig. 7a**). Since it is straightforward to create concave and convex patterns by micromolding (**Fig. 5b**), we tested whether the direction of the fractal topography affected podocyte cultivation using the Mosaic assay. Interestingly, cell coverage was significantly higher on convex versus concave patterns (**Fig. 5c-e**) despite comparable cell morphology Df (**Fig. 5f**).

We were able to fabricate fractal well plates with customized topographical bottoms covered with high density HF patterns and confirmed successful human podocyte culture with Matrigel coating (**Supplementary Fig. 8**). We then compared podocyte culture using this fractal topographical well plate versus a standard flat tissue culture well plate. Mouse podocytes exhibit a higher gene expression of *Nphs1* upon differentiation on the fractal plate (**Fig. 8h**). Mosaic human podocytes show a more in vivo-like morphology, as reflected by a higher Df and a lower aspect ratio (**Fig. 8j-k**). We also performed an injury study induced by puromycin aminonucleoside (PAN) as proof-of-concept for the use of this fractal plate for drug testing (**Fig. 8i**). Upon PAN injury, foot process effacement shown by a decrease in cell morphology Df was observed as expected in Mosaic human podocytes cultivated on both the standard and fractal well plates. Cells on the fractal plate appear to be more sensitive with the Df of their morphology decreasing to a greater extent (**Fig. 8j**).

### 2.6. Distinguishable gene expression profiles indicate enhanced ECM deposition, remodelling and signal transduction on fractal topographical substrates

To explore the changes of cellular transcriptome that possibly drove the effect we observed on fractal topography, we analyzed published RNA sequencing data^32^ from primary fetal human podocytes, which is a more authentic cell source than cell lines, grown in a previous work on identical NT, RT, and HF substrates^13–15^ as well as cells prior to culture considered as the baseline. Importantly, our samples here were cultivated on identical substrates as those from the previous bulk RNA sequencing work ^13,15^. The data are published and available on GEO repository under accession code GSE185491,^32^ where frozen indicates baseline cells. Not only did cells cultured on the three substrate groups effectively form distinct clusters among themselves and against baseline cells by unsupervised principal component analysis (PCA) (**Fig. 6a**), the Keygenes analyses (**Fig. 6d-e**), also confirmed a more mature age and a more definitive kidney identity of cells on fractal substrates and of Yoshimura *et al*.^6^ datasets in comparison to the baseline cells, which is further supported by the PluriTest analyses. PluriTest was able to demonstrate a shift towards maturation upon fractal shape stimulation when compared to published signatures^6^ and classify both biochemically and topographically stimulated cells as more mature than nephron progenitor cells, with biochemical iPSC-podocytes reaching a higher degree of maturation on the PluriTest score (**Fig. 6b**). Improvements in the topographical groups (HF and RT) vs. flat with respect to maturation assessed by PluriTest in comparison to the baseline cells were also noted (**Fig. 6c**).

**Fig. 6.**
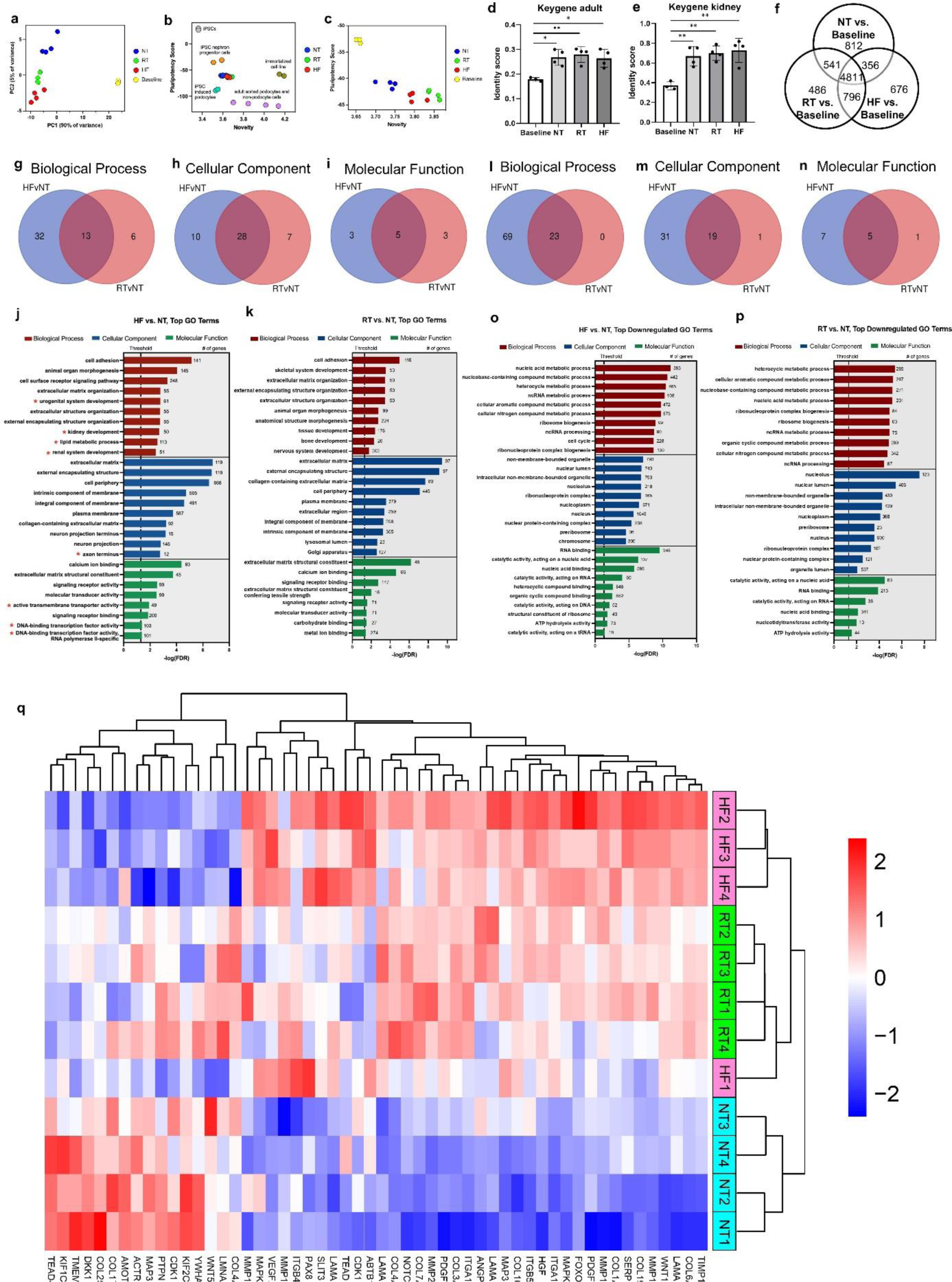
RNA sequencing suggests that fractal topography modulates human podocyte response via ECM deposition, cell adhesion and kidney development. **a**, Unsupervised principal component analysis (PCA) delineating clusters in total gene expression patterns for four main conditions: baseline, NT, RT, and HF. **b,** Pluripotency scores compared to published datasets of undifferentiated induced pluripotent stem cells (iPSCs), iPSC-derived progeny, adult podocytes and an immortalized cell line^6^. **c,** PluriTest ran separately on baseline, NT, RT and HF samples. **d-e,** KeyGenes analysis comparing baseline cells (frozen) and different culture conditions (NT, RT and NF) to the **d,** adult stage of development and **e,** kidney identity^6^. Data plotted as mean ± SD. One-way ANOVA with Tukey’s post-hoc test, *p≤0.05, **p<0.01. **f,** Venn diagram of differentially expressed genes in the three culture conditions (NT, RT and HF) compared to the baseline cells, prior to cultivation. **g-i**, Venn diagrams illustrating the number of upregulated overlapping and distinct gene ontologies (GO) terms for **g,** biological processes, **h,** cellular components and **i,** molecular function. **j-k,** Top 10 most upregulated GO terms for biological processes, cellular components and molecular function in **j,** HF vs NT and **k,** RT vs NT. *GO terms exclusive to HF vs. NT and not significant in RT vs. NT. **l-p**, Venn diagrams illustrating the number of downregulated overlapping and distinct gene ontologies (GO) terms for **l,** biological processes, **m,** cellular components and **n,** molecular function. **o-p,** Top 10 most downregulated GO terms for biological processes, cellular components and molecular function in **o,** HF vs NT and **p,** RT vs NT. **q,** Supervised hierarchical heatmap of ECM, matrix remodelling, cell adhesion and signaling genes relevant for podocyte function with significant differential expression. RNA sequencing data from primary GW18 fetal human podocytes after 3 days in culture on n=4 scaffold samples per group and baseline cells prior to culture available on GEO repository under accession code GSE185491 was used for analysis.^32^ NT, no topography; RT, round topography; LF, low fractal; HF, high fractal.

When examining the differences in gene expression with respect to baseline cells, majority of differentially expressed genes were found to be common to all the groups (4811 genes, **Fig. 6f**) as expected upon cell thawing. Baseline cells were substantially different from cultivated cells with additional differences emerging with topographical culture (i.e. unique 486 genes on RT and 676 on HF, **Fig. 6f**). Unsupervised k-means clustering determined eight clusters for which gene set enrichment analysis identified enhanced signaling pathways **(Supplementary Fig. 9a)**. k-means cluster analysis for biological process and KEGG suggested the dose response with topographical cues, where the HF group was most distinct from the baseline cell group (**Supplementary Fig. 9b-c**). In a dose-like manner, HF had enrichment in pathways associated with extracellular structure and matrix organization, nephron and epithelial development, motility, and structure formation. The dip in Cluster 4 on NT, may be related to de-differentiation on flat substrates, as gene ontologies (GO) for biological process included ECM organization and KEGG for cell adhesion molecules were identified amongst important processes and pathways that were downregulated in NT with respect to baseline cells (**Supplementary Fig. 9c**). In contrast, Cluster 5 demonstrated highest gene expression on HF in important processes such as collagen activated signaling, ECM organization and extracellular structure organization (**Supplementary Fig. 9b**).

Next, we looked at most significantly enriched gene ontology (GO) terms among all genes that were significantly upregulated in HF vs. NT and RT vs. HT. The results indicated that topography in general resulted in upregulated cell adhesion and organ morphogenesis biological processes as well as upregulated ECM, cell periphery and plasma membrane cellular components (**Fig. 6g-k**). Yet, when HF vs. NT was contrasted to RT vs. NT, only HF group exhibited upregulation in GO terms for kidney development, renal system development, regulation of anion transport, urogenital system development, cell communication, metanephros development, cell-cell adhesion, vesicle mediated transport and cell-cell signaling (red stars in **Fig. 6j, Supplementary Data 1** and **2**). In terms of molecular function, DNA binding transcription factor activity was uniquely upregulated in HF vs NT, in comparison to RT vs NT (**Fig. 6j**). On the other hand, the top 10 most significantly downregulated genes in the HF group compared to other groups, were related to metabolic process and cell cycle (**Fig. 6l-p**), consistent with low proliferation levels known as the effect of topographical cues.^33^ Notably, neuron projection and axon terminus were found to be the top 10 most upregulated GOs in the HF group (**Fig. 6j**). Given the peculiar similarity in terms of cellular morphology between polarized neurons and podocytes (prominent cell body, polarized protrusion network), commonalities between structural organization of podocytes and neurons have been previously recognized.^34,35^

In addition, we examined expression of several ECM molecules, integrin receptor subunits, mitogen activated kinase (MAPK) isoforms, matrix metalloproteinases (MMPs) and signal transduction molecules that exhibited significant differences between the HF group vs. NT alone or between both HF vs. NT and HF vs. RT (**Fig. 6q**). Clearly, topographical substrates exhibited a significantly higher expression of key collagen and laminin isoforms (*COL6A3, COL1A2, COL19A, COL16A1, LAMA2, COL3A1, COL7A1, COL4A5, LAMA5, LAMA3*) with notably high expression in the HF condition, whereas *COL17A*, reported to be implicated in the growth of multi-layered transformed epithelium was downregulated in HF group.^36^ *COL4A5* and *LAMA* subunits are key structural components of the glomerular basement membrane. *COL4A5* in particular, is one of the critical determinants of glomerular basement membrane structural integrity and aberrant expression of this protein is linked to numerous glomerular kidney diseases, e.g. focal segmental glomerulosclerosis (FSGS), Alport’s, diabetic nephropathy, etc. Key matrix remodelling enzymes were also upregulated on topography and generally higher in the HF group (*TIMP1, MMP13, MMP14, MMP15, MMP16*) along with integrin receptor subunits (*ITGA10, ITGB5, ITGA11*).

*PAX8* which regulates branching morphogenesis was higher on HF substrates, and so was *SLIT3*, whose knockdown results in defective kidney development.^37,38^ Differences in gene expression of important signal transduction proteins (e.g. *NOTCH3*, high in HF) and morphogens (e.g. *WNT5A* and *DKK1* low in HF), transcription factors (e.g. YAP pathway molecule *AMOTL2*, low in HF) and kinases (*MAPK3* significantly higher in both HF vs. NT and HF vs. RT) of relevance to glomerulus development and kidney function^39,40^ presented with possible candidates for elucidation of biochemical drivers of the observed branching behaviour (**Fig. 6q**). These data also solidify the utility of the topographical substrates, particularly HF substrate, as important determinants of podocyte maturity and differentiation. Interestingly, the expression of angiogenic factors (*VEGF, PDGFA, HGF, ANGPT1*) was enhanced on topographical substrates, which may be important in future tissue engineering applications (**Fig. 6q**).

Lastly, we identified a subset of genes in the categories relevant for the podocyte and kidney function that were differentially expressed between HF and RT group, but not between the HF and NT group (**Supplementary Fig. 10**). Intriguingly, these genes, which were specifically higher in RT than in HF were reported to be upregulated in cancers in general and more specifically in renal cancers (*COL12A1,*^41,42^ *COL5A1,*^43^ *COL5A2*^44^ and *MMP17*^45^) and other kidney diseases such as: acute kidney injury (*MAP3K14*^46^), focal segmental glomerulosclerosis (*COL4A2*^47^), renal fibrosis (*ITGAV*^48,49^) and polycystic kidney disease (*ITGB1*^50^). This indicates that although topography generally results in the upregulation of ECM, cell-matrix attachment and cytoskeletal re-arrangement genes in comparison to the flat substrates, a more physiologically relevant topography such as HF, prevents overactivation of genes that are often linked to pathological conditions such as cancer. This further reinforces the importance of fractality as an ideal dose of disorder required for physiological function in many systems.^4,51^

### 2.7. YAP signaling pathway plays a role in maintaining podocyte morphology

Upon searching the published RNA sequencing data^32^ from human podocytes grown on fractal substrates, we identified key members of YAP1 interactome (generated using GeneMANIA algorithm^52,53^) that were differentially regulated on the HF substrates in comparison to NT and RT (**Fig. 7a-b**). YAP interactome members were found to be differentially regulated on HF substrates (**Fig. 7a**).

**Fig. 7.**
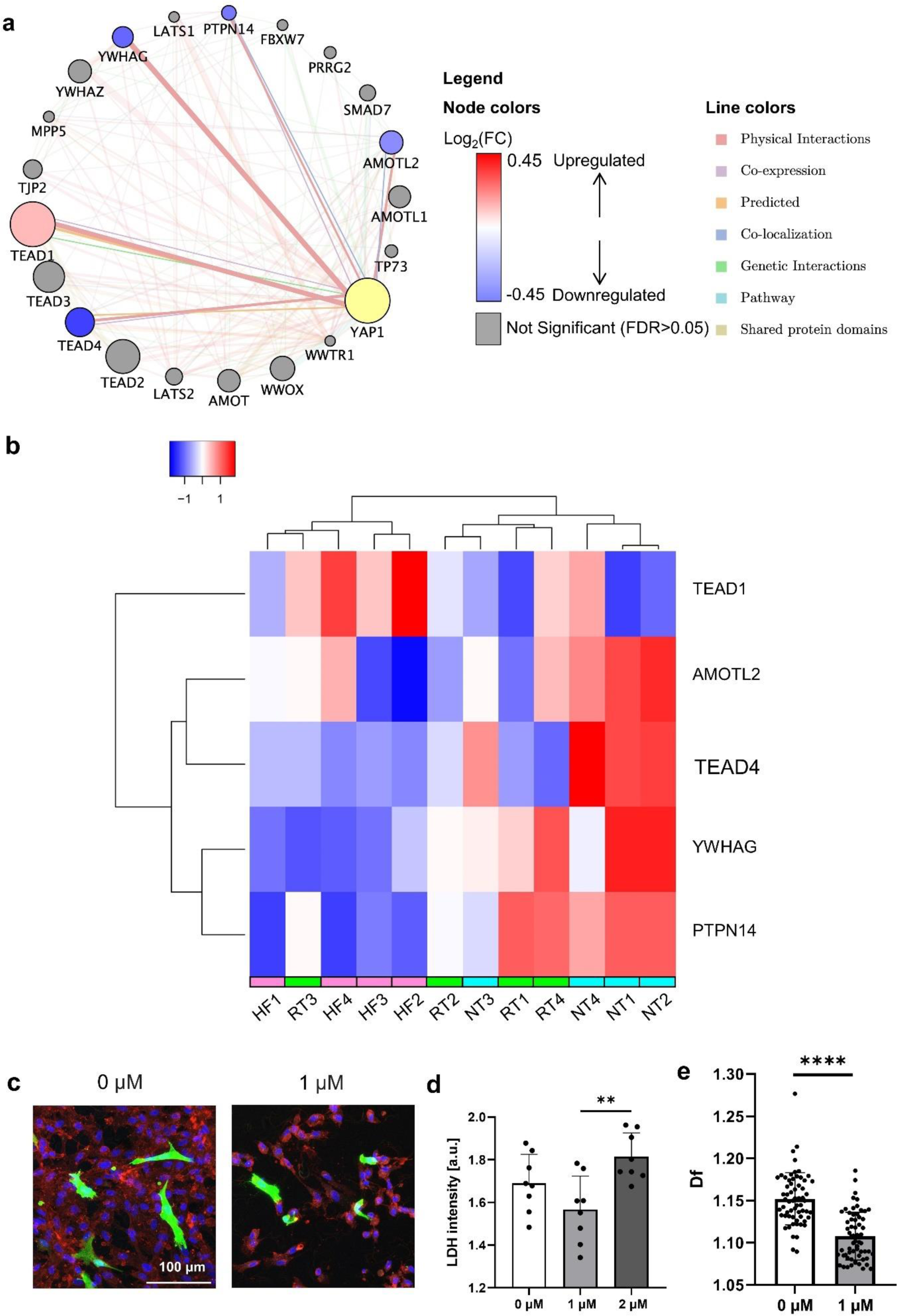
YAP signaling pathway plays a role in maintaining human podocyte morphology on fractal substrates. RNA sequencing data from primary GW18 fetal human podocytes after 3 days in culture on n=4 scaffold samples per group and baseline cells prior to culture available on GEO repository under accession code GSE185491 was used for analysis for **a** and **b**.^32^ **a**, YAP interactome. The nodes represent the related genes identified by GeneMANIA, with the size of each node indicating the rank of how closely the gene was associated with YAP1. The node colours are used to relate our RNA seq data to the YAP1 interactome. Blue nodes are genes of the YAP1 interactome that are downregulated in HF vs. NT. Red nodes are genes of the YAP1 interactome that are upregulated in HF vs. NT. Grey nodes are still part of the YAP1 interactome according the GeneMANIA algorithm, but had no significant difference between HF and NT from the RNA seq data. The lines connecting nodes describe the types of interactions between the nodes. **b,** Heatmap of significantly different genes from YAP interactome. **c,** Mosaic human podocyte assay on HF scaffolds with and without treatment using YAP inhibitor verteporfin. **d,** Lactate dehydrogenase (LDH) release of Mosaic assay cells inhibited with verteporfin at 0 µM, 1 µM, and 2 µM. n=8 scaffold samples per group. One-way ANOVA statistical testing with Tukey’s test for multiple comparisons. **e,** Quantification of whole cell morphology by fractal dimension of Mosaic assay cells on HF scaffolds inhibited with verteporfin. n=3 samples per group. Student t-test used for statistical testing. **p≤0.01, ****p<0.0001. Data plotted as mean ± SD. **a-b**, primary fetal human podocytes. **c-e**, immortalized human podocyte cell line. NT, no topography; RT, round topography; LF, low fractal; HF, high fractal.

*TEAD1*^54^ which is primarily responsible for cell adhesion, and motility was upregulated in HF substrates, whereas *TEAD4* which is known to supress collagen 1 promoter activity^55^ was downregulated on HF consistent with enhanced matrix deposition and remodelling in this group (**Fig. 7a-b**). *YWHAG* which is primarily responsible for proliferation was downregulated on HF, consistent with its effect of reduced proliferation. AMOT family (downregulated *AMOTL2* in HF), inhibits *YAP1* by regulating its localization and promoting its phosphorylation,^56^ while *PTPN14* which was downregulated in HF, regulates YAP intracellular localization and inhibits its transcriptional co-activator activity.^57^ These findings suggest that *YAP1* activity may be responsible for the observed effects of cellular branching.

Upon YAP inhibition in human podocyte cell line on HF substrates (**Fig. 7c**) in the presence of suppressor of YAP–TEAD complex (Verteporfin^58^) at the dose of 1μM, which did not show any significant deterioration of cell viability (**Fig. 7d**), Df of podocyte morphology was decreased significantly in comparison to the inhibitor-free control (**Fig. 7e**). Collectively, these results support the involvement of YAP in cell shape control on fractal substrates.

## 3. Discussion

We highlight here the fractal nature of glomerular podocytes and the glomerulus itself, that may ultimately prove to have significant contribution to the functional identification of healthy vs. diseased states. Our findings suggest how fractal topographical cues could enhance cell maturation: fractal topography organizes ECM proteins and facilitates localization and spatial arrangement of subcellular structures, resulting in higher-order branching morphology and polarity. We confirmed this behaviour by using or considering three cell sources: conditionally immortalized mouse podocyte line, non-conditionally immortalized human podocyte line, and published RNA sequencing data^32^ from primary human fetal podocytes, as each source offers differences in terms of proliferation rate, genetic fidelity, and physiological relevance that are important for the validation process. Each cell source has strengths and weaknesses.

The mouse cell line was conditionally immortalized, thus it has a thermoswitching mechanism that could deactivate the forced cell cycle processes and slow proliferation and allow differentiation over longer cultivation periods, namely, of at least 14 days. The non-conditionally immortalized human cells do not have a mechanism built-in for switching off the genetically induced immortalization. These cells keep proliferating past confluence and this limits the ability to achieve a fully differentiated phenotype. While they represent a human source, their cultivation period is short before cells begin overcrowding. On topographical substrates, the cultures could only last ∼5 days before overgrowing and becoming prone to being peeled off from the PDMS substrates during handling of the scaffolds.

The primary cells are the most authentic and unmodified cell source, however, limited in number and more immature by definition as they are fetal-derived. In this case, the published RNA sequencing data used for analysis were generated from podocytes from gestation week 18 via a commercial source (Lonza) ^13–15^ ^32^. At this stage of development, podocytes are still relatively immature and able to undergo proliferation as they are still developing and populating the developing nephron and glomerulus.

In their native environments, cells and tissues exhibit complex fractal configurations, an attribute scarcely replicated in traditional in vitro cultures. The impact of mimicking these in vivo fractal patterns on cellular development and response to structural stimuli remains underexplored. Introducing fractal dimensions as an external variable may be pivotal for augmenting in vitro cell differentiation. We emulated the fractal designs found in glomerular histology to fabricate three-dimensional (3D) substrates for cell culture. Our work captured some of the fractal effects by fitting a single power law to morphologies of podocytes and glomeruli that are in reality multi-fractal. Future work should build upon this concept by breaking down the multi-fractality of the system and more precisely incorporating it into experimental design. Furthermore, building upon the 2.5-D fractal substrates, fractal scaffolds in a full 3-D configuration and at a physiologically relevant scale would further approximate podocyte microenvironment and consequently bring their maturation level closer to the in vivo state.

YAP activation, through nuclear localization is required for podocyte function,^59^ as well as nephron development, as conditional knockdown of YAP in mouse kidney led to reduced nephrogenesis and morphogenesis, in a manner that was independent of proliferation and apoptosis,^60^ consistent with our observation of YAP1 involvement in shape patterning.

In summary, podocytes cultured on the fractal matrices demonstrated elevated levels of functional biomarkers and more sophisticated cellular architecture. RNA sequencing indicated that podocytes on fractal substrates show enhanced extracellular matrix (ECM) synthesis and alteration, driven by the YAP signaling pathway, in contrast to counterparts cultured on flat or non-fractal surfaces. Our integration of fractal topography into standard tissue culture plates could offer a straightforward and effective approach for more precise cell culture.

## 4. Methods

*Extraction of Histological Features of the Glomerulus*: Kidney histology images from the literature were imported to Fiji and cropped to select the glomerular region. Next, the images were thresholded using the Otsu filter in Fiji to create a binary mask that cover the tissue sections, followed by despeckling once to reduce the background noise. The outline of the binary mask was then collected in Fiji for fractal dimension analysis.

*Fractal Dimension Analysis*: The fractal dimension analysis was performed using either the box counting algorithm from the Fraclac plugin in Fiji or the box-counting method through a Matlab code developed by Frederic Moisy.^61^ Fraclac was used to generate single-value Df; the Matlab code was used for multi-fractal analysis. Briefly, outlined binary images were input to Fraclac and box counting with the white background selected was applied to calculate fractal dimension. The method developed by Frederic Moisy involves covering the image with a set of boxes of size R, and counting the number of boxes (N) that are required to cover the entire image. The relationship between N and R can be expressed as:

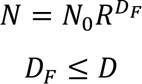

where D_F_ is the fractal dimension or the Minkowski-Bouligand dimension, and D is the space dimension. By plotting the local exponent, we can identify the local fractal properties within a limited range of box size R using the equation:

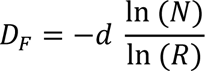

*Fast Fourier Transform (FFT) Analysis*: We employed the FFT analysis to investigate the frequency components present within our images. The FFT function available in ImageJ was used to compute the FFT image of our data. The resulting FFT image was then subjected to surface plotting and profile plotting of the central vertical line.

*Topographical Pattern Fabrication*: The design of the topographical patterns RT, LF, and HF is reported previously.^13–15^ The ST pattern was designed in AutoCAD to approximate the sharp and pointy features of the HF pattern by arranging lattices of spikes (circles of 10 μm diameter, 12.5 μm between centers of adjacent circles) within a circular region (150 μm in diameter) in close proximity to the size of a repeating unit of the HF pattern). The ST unit design, packed hexagonally similar to the HF pattern, was imported to a direct laser writer (MicroWriter ML3 Baby) and replicated to span a square large enough to cover the entire area of a well in a standard tissue culture 24-well plate. The master molds with topographical patterns were fabricated using a method developed and reported previously.^13–15^ Briefly, a silicon wafer was spin coated with HMDS at 3000 rpm for 30 seconds, followed by softbaking at 150 °C for 1 minute, to increase the adhesion of the subsequent photoresist layer. Positive photoresist AZ P4620 was then spin coated on top of the HMDS layer at 1500 rpm for 30 seconds, followed by softbaking at 110 °C for 80 seconds and rehydrating for approximately 30 minutes at room temperature. The design patterns were written onto the wafer by exposing the photoresist to laser at 770 mJ/cm^2^, followed by developing and reflow (120°C for 1-2 minutes) to smooth the edges of the features.

*PDMS Substrate Fabrication*: Patterns from master molds fabricated by direct laser writing of AZ P4620 positive photoresist on silicon wafer were copied to PDMS negative molds by inverse molding at a ratio of 1:10. The PDMS negative molds were then fully cured by baking at 120°C overnight and used as master molds to fabricate PDMS substrates used for cell culture. Circular PDMS substrates that fit in standard tissue culture well plates were hole punched using a biopsy punch.

*Substrate Pattern Characterization*: Patterned substrates were imaged with a 3D digital microscope (Keyence) in z-stacks to acquire their surface profiles. The stacked microscope images were then reconstituted into 3D display. Patterned substrates were also imaged under scanning electron microscopy (SEM, Hitachi TM4000 or SU7000) at 5.0 kV voltage acceleration to confirm feature transfer from master molds to substrates.

*Podocyte Culture*: The conditionally immortalized murine podocyte E11 cell line was a generous gift from GSK. Murine podocyte E11 cell line was expanded in flasks coated with 0.1 mg/ml rat tail collagen I (Corning, #354236) under the proliferative condition at 33°C with 10 units/mL mouse interferon gamma supplemented in fresh in RPMI 1640 basal media containing HEPES and GlutaMAX (Gibco, #72400120) with 10% FBS and 1% pen-strep. To induce differentiation, cells were cultured under the differentiation condition at 37 or 38°C in DMEM-F12 basal media with 10% FBS and 1% pen-strep. The immortalized Podo/Tert256 human podocyte hTERT cell line (Evercyte) was expanded in flasks coated with 6 µg/cm^2^ human collagen I (Sigma, C7774) at 37°C in MCDB131 basal media (Gibco, #10372019) supplemented with 20% FBS, 2 mM GlutaMAX (Gibco, #35050061), 12 µg/mL Bovine Brain Extract (Lonza, CC-4098), 10 ng/mL hEGF (Sigma-Aldrich, E9644), 25 ng/mL hydrocortisone (Sigma-Aldrich, Cat# H0396), and 100 µg/mL G418 (InvivoGen, ant-gn-5). For both mouse and human podocytes, media was changed every 2-3 days.

*Substrate Preparation for Cell Seeding*: PDMS substrates were sterilized by autoclave and inserted in well plates to allow for standard cell culture techniques to be applied for cell seeding and cultivation on the scaffolds. In addition to scaffolds, a blunt tweezer was autoclaved at the same time and then used to insert the scaffolds into the wells of a tissue culture well plate. Each scaffold was pushed to the bottom of the well using the tweezer to ensure all edges of the scaffold were flat in the well and to eliminate the gap space between the scaffold and the well. By doing so, fewer cells would be lost in the gap space and not settle on the substrate surface when the cells were seeded. After inserting the scaffolds, PBS was added to the wells to wet the scaffold surface and the well plate was incubated at 37°C for at least 30 minutes. Prior to cell seeding, ECM proteins were coated on top of the substrates to ensure adequate cell adhesion. Briefly, Matrigel (Corning, #354234) diluted 1:60 was coated on substrates used for mouse podocyte cultivation. Matrigel (Corning, #354234) diluted 1:60 or porcine kidney ECM (Xylyx Bio, NativeCoat Kidney ECM surface coating kit #MTSKY201) diluted according to manufacturer’s instructions at 0.08 mg/mL was coated on substrates used for human podocyte cultivation. Appropriate coating is critical for successful cell culture, and natural ECMs such as porcine kidney may exhibit batch-to-batch variability that will require optimization of the coating procedure and cell seeding density. Most consistent and reproducible results are obtained if Matrigel is used for coating.

*Cell Seeding*: Mouse podocytes were seeded at 50,000 cells/cm^2^ and allowed to adhere and reach ∼80% to full confluence under the proliferative condition before thermoswitching and culturing under the differentiation condition as described in the Podocyte Culture section. For the early study endpoint (immature cells), mouse podocytes were cultivated under the differentiation condition for 3 days. For the late study endpoint (mature cells), mouse podocytes were cultivated under the differentiation condition for 14 days. Human podocytes were seeded at 50,000, 100,000, or 200,000 cells/cm^2^, and grown until ∼80% to full confluence as the study endpoint (3-5 days). To seed the cells, harvested cells from the expansion cultures were resuspended at a concentration corresponding to the target seeding density. After that, the cell suspension was pipetted up and down a few times to mix evenly and then added to each well. Finally, the well plate was left in the incubator (37°C for human podocytes, 33°C for mouse podocytes) for the cells to attach. Media change was performed the next day or in two days. The same culture media was used for human podocytes whereas media was switched to differentiation media for mouse podocytes.

*Glycocalyx Imaging*: For glycocalyx imaging, mouse podocytes were fixed with 4% PFA, 1% glutaraldehyde, and 1% Alcian Blue overnight. Next, the samples were post-fixed with 0.5% osmium and dehydrated in ethanol in a serial manner, followed by critical point drying (CO_2_) and gold coating. Samples were imaged using a Hitachi SU-8230 scanning electron microscope at 2.0 kV voltage acceleration.

*Immunofluorescent Staining*: At the study endpoint, cells were fixed with 2% or 4% PFA for 20 minutes at room temperature and rinsed with PBS for three times. Next, cells were blocked in 5% normal goat serum (NGS) with 0.1% triton X-100 for 1 hour under gentle shaking. Without rinsing, cells were incubated with primary antibodies diluted in 2% NGS with 0.04% triton X-100 overnight at 4°C. After that, cells were washed with PBS for three times, followed by incubating with secondary antibodies and stains in 2% NGS with 0.04% triton X-100 for 1 hour at room temperature under gentle shaking. Samples were then washed three times with PBS and carefully taken out of the wells. Excess liquid was removed from the bottom of each scaffold with paper towel. Two to three scaffolds were placed on each microscope slide. 6 µL of DAPI-containing mounting media (Vector Laboratories, Vectashield #H-1200) was added to the surface of each scaffold. Lastly, a cover slip was placed on top of each scaffold to spread the mounting media over the entire surface of a scafold. Care should be taken when handling a scaffold to make sure not to disturb the delicate sheet of cells on top. When taking a scaffold out of the well, efforts should be made to minimize any contact with the cell side of the scaffold and to avoid twisting or bending the scaffold during handling. During the staining procedure, scaffolds should be rinsed gently with solutions gently added from the side as opposed to on top. The antibodies and fluorescent stains used are listed in **Supplementary Table 1**.

*Immunofluorescent Imaging and Analysis*: Samples were imaged via confocal microscopy using a Leica Lightsheet or Nikon A1R confocal microscope. Z-stacked images were taken to capture cell morphology on topographic substrates. Maximum intensity projection images were acquired from the z-stacks for image analysis. Regions of interest in the images from the experiment comparing the HF group with the ST control were selected in Fiji as circular areas containing the patterns. To quantify fluorescence signals, the channel of interest was thresholded by filter in Fiji and integrated and mean intensity were measured. Directionality analysis was performed using MATLAB to compute eccentricity and orientation variance of the signals from subcellular structures. Eccentricity is defined as a measurement of the elliptical shape of structures where a score of 0 is considered a circle and a score approaching 1 indicates elongation of structures. Briefly, the images were converted to a binary format and the **imageprops** function was used to measure eccentricity and orientation. The averaged eccentricity value of all elements normalized to the ST control was reported. The orientation variance within each image normalized to the ST control was presented. The images from the experiment with four substrates (NT, RT, LF, HF) were analyzed as a whole without selecting regions of interest. Integrated intensity of the fluorescence signals from these images were quantified in Fiji as described above, averaged by cell numbers, and reported as values normalized to the average value from the day 3 NT group. The same settings were applied to all the samples during image acquisition, processing and analysis.

*Mosaic Assay*: Mosaic Podo/Tert256 human podocytes (a mixture of GFP-labelled cells and non-fluorescent cells at a ratio of 1:11) were cultured and seeded in the same way as regular human podocytes as described above. At the study endpoint, Mosaic cells were fixed with PFA, followed by staining with rhodamine-labelled wheat germ agglutinin (WGA) and mounting with DAPI-containing mounting media as described above. Samples were imaged via confocal microscopy using a Zeiss LSM 880 Super Resolution or Leica Lightsheet Confocal microscope. Z-stacked images spanning the entire section with visible signals were taken for each sample. The images were processed using the ZenDesk software to increase the contrast of GFP-positive cells’ morphology before cell selection in Photoshop by setting gamma in the FITC channel to 2.5 with the white point adjusted to 30 and adjusting the white point of the DAPI channel to 80. In Photoshop or Fiji, the wand tool was used to select fluorescent cells from which binary masks of single cells were created and exported. Single cells were verified by overlaying the DAPI channel with the FITC channel to ensure each cell only contained one nucleus. The outline of each binary images was obtained in Fiji, and fractal dimension was calculated using the FracLac plugin as described above.

*EV Isolation*: At the study endpoint, mouse podocytes were washed with PBS twice and cultured in serum-free culture media for a period of 48 hours. Media from 8 scaffolds for each topographical group was combined and collected in a 15 mL centrifuge tube. Cell debris were removed by centrifuging at 3000 g for 10 minutes at 20°C. EVs were then isolated by the miRCURY Exosome Isolation Kit (Qiagen, #76743) according to the manufacturer’s protocol. Precipitation buffer was applied at a ratio of 4:10 to the samples, followed by rotating overnight at 4°C for the precipitation of EVs. Precipitated EVs were then centrifuged at 3200 g for 30 minutes at 20°C. After removing supernatant, the pellet was gently rinsed with PBS. 100 µL resuspension buffer was added to isolated EVs prior to downstream analysis.

*Western Blotting*: Mouse podocytes were differentiated in 24-well plates, with 6 wells per plate dedicated to each of the 4 surface topographies (NT, RT, LF, HF). On day 14 of culture, cells were washed with ice-cold PBS. 150μl of RIPA buffer supplemented with a protease inhibitor tablet (Thermo Scientific) was added to a single well for podocyte lysis, pipetting up and down several times while scraping the surface of the well with the pipette tip to release cells. The full volume of RIPA-lysate mixture was then transferred sequentially to the next 5 wells of like-topography one at a time, repeating the process of pipetting and scraping each time. After all 6 wells of a single topography were lysed, the RIPA-lysate mixture was transferred to a microcentrifuge tube and lysis was repeated for the remaining 3 topographies.

EVs were isolated from conditioned media of day 14 cultures of mouse podocytes grown on NT and HF topographies as previously described. EV pellets were lysed in 50μl RIPA buffer supplemented with protease inhibitor. Cell and EV lysates were centrifuged at 10,000g and 4°C for 15 minutes. Supernatants were transferred to fresh tubes and pellets were discarded. The Pierce BCA Protein Assay Kit (Thermo-Fisher Scientific) was used to analyze protein concentration in lysates. Initial western blots were performed to determine the linear regime for sample loading and target detection. For podocyte cell lysate western blots, samples were loaded at 7µg total protein adjusted to a volume of 35μl per well of a 15-well gel; EV blots were loaded at 10µg total protein per adjusted to a volume of 40μl per well of a 10-well gel.

Appropriate sample volumes were diluted with 4x Sample Loading Buffer (LI-COR), 10x BOLT Sample Reducing Agent (Thermo Fisher Scientific), and DI water, then incubated at 70°C for 10 minutes. Samples were loaded into Bolt 4-12% Bis-Tris Gels (Invitrogen). Electrophoresis was perfomed in Bolt MOPS SDS Running Buffer (Invitrogen) at 200V for 35 minutes or until protein front reached the bottom of gels. Completed gels were rinsed in DI water then transferred to Immobilon-FL PVDF membranes (Millipore Sigma) at 20-25V for 7 minutes using the iBlot Gel Transfer Device and transfer stacks (Invitrogen).

REVERT Total Protein Stain (LI-COR) was applied to PVDF membranes according to the manufacturers protocol before imaging on the LI-COR Odyssey Fc Imager (LI-COR). Blocking was then performed by incubating membranes in skim milk buffer for 1 hour. Membranes were washed and stained overnight at 4°C with anti-PODXL (Invitrogen, PA5-28116, 1:1000 concentration), anti-CD63 (Abcam, ab217345, 1:1000 concentration), or anti-ALIX (Thermo-Fisher Scientific, MA5-32773, 1:1000 concentration). Membranes were then washed, appropriate IRDye800CW secondary antibodies were added (LI-COR, 1:10,000 concentration) for 2 hours, washing was repeated, and images were captured with a 10 minute exposure. PODXL blot was stripped using a mild stripping buffer consisting of 1.5% w/v glycine, 0.1% w/v SDS, and 1% v/v tween 20, titrated to pH 2.2 in deionized water. Blot was incubated in stripping buffer twice for 5 min, washed four times, then blocked as previously described. PODXL blot was then washed and stained as before using anti-GAPDH (Invitrogen, PA1-987, 1:1000 concentration) and IRDye800CW secondary antibodies prior to imaging.

*Transmission Electron Microscopy of EVs*: Plasma treatment was used to pre-treat electron microscopy grids (Electron Microscopy Sciences). Grids were incubated with 5µl of 100x EV concentrate for 1 minute before wicking with a wet filter paper. To wash grids, 5µl of DI water was applied then wicked away. 5µl of 2% uranyl acetate was then added to grids for 30 seconds as a negative stain. Grids were once again wicked and imaging was subsequently performed on a Talos L120C TEM (Thermo-Fisher Scientific) at 22,000x, 45,000x, and 92,000x.

*YAP Inhibition Study*: HF and NT scaffolds were prepared as described above. Mosaic human podocytes were seeded on top of the scaffolds at a concentration of 50,000 cells/cm^2^ (100,000 cells per scaffold). Verteporfin (1 or 2 μg/ml, Tocris) was added to the culture media 24 hours post seeding. Samples were fixed on day 3. Mosaic assay imaging and analysis were performed as described above. To validate cell viability, lactate dehydrogenase assay (LDH) assay (Cayman) was performed on media collected on day 5 of culture according to manufacturer’s protocol.

*Fractal Topographic Well Plate Fabrication*: To produce 24-well plates with a high density fractal pattern, polystyrene sheets were hot embossed against a master mold designed and laser patterned with high density fractal units. The polystyrene base was then cut to the size of a 24-well plate bottom and bonded to the bottomless well plate using PDMS as a glue. To produce 384-well plates with a high density fractal pattern, extruded, thermoplastic elastomer sheets (230μm thick, Vertex FK79, Gel-Pak Inc, USA) were hot embossed for 10 minutes at 160°C (EVG520HE, EV Group, Austria). The base was then sealed to a self-adhesive bottomless well plate from Grace Bio-Labs. 96-well plates with a high density fractal pattern and TPE bottoms were fabricated in a similar method using a master mold designed for a 96-well blueprint.

*PAN injury model:* Mosaic human podocytes were seeded on the standard and fractal 96-well plates at a concentration of 50,000 cells/cm^2^. PAN (10 μg/ml, Sigma-Aldrich, P7130) was added to the culture media 24 hours post seeding. Fresh media containing 10 μg/ml PAN was changed on day 3. Cells were fixed on day 5. Mosaic assay imaging and analysis were performed as described above.

*RNA Isolation*: RNA extraction and isolation of mouse podocytes was performed using the PicoPure RNA isolation Kit (Thermo Fisher Scientific, #KIT0204) according to manufacturer’s protocol (100 µL instead of 50 µL of lysis buffer was applied to extract RNA), including DNase treatment with RNase-Free DNase Set (Qiagen, #79254). At the final step, 12 µL of elution buffer was used. The concentration and quality of isolated RNA was measured using a NanoDrop spectrophotometer. Isolated RNA was stored at -80°C.

*Quantitative polymerase chain reaction*: Isolated RNA was reverse transcribed into cDNA using the SuperScript™ II RT kit (Invitrogen, #18064-014) according to the manufacturer’s protocol. Briefly, random hexamer primer (ThermoFisher, #SO142) was mixed with total RNA (300 ng), dNTP mix (Invitrogen, #18427-013) and water according to the specified quantities. The mixture was heated to 65°C for 5min then chilled on ice. Each of the remaining contents of the SuperScript™ II RT kit were then added and incubated as specified in the protocol, with the addition of 1 µL of RNaseOUT™ (Invitrogen, 10777-019) to each preparation. The RT mixture was incubated at 25°C for 10min, 42°C for 50min, and 70°C for 15min. Each final 20 µL cDNA prep was diluted 1:3 in RNase-free sterile water. qPCR was carried out following the PowerTrack™ SYBR™ Green Master Mix (ThermoFisher, #A46109) kit protocol and performed on a CFX384 Touch Real-Time PCR Detection System. A 10 µL reaction volume was used for each PCR well in a 384 well plate, adding diluted cDNA, yellow sample buffer, master mix, and water as specified in the kit protocol. A list of forward and reverse primers used is listed in **Supplementary Table 2**. For comparison of gene expression, analysis was performed using the ΔΔC_t_ protocol, with GAPDH as the reference gene and averaged ΔC_t_ values for NT as the control condition from which to determine ΔΔC_t_ differences and fold changes for NT and HF data points.

*KeyGenes Analysis*: Data were analyzed from the the GEO database, accession number GSE185491. In order to quantify the tissue type and the developmental stage of the differentiated podocytes at the different topographical states, we used the identity score (range 0-1) from the KeyGenes tool.^62^ The identity score predicts the “identity” of a test sample to a known feature (tissue or age) by comparing gene expression profiles of classifier genes used in KeyGenes. A higher predicted identity score implicates a greater identity to a specific tissue or differentiation age. The identity score for the topographic replicates against kidney tissue, and adult stage of development is plotted as barplots (mean ± SD) using GraphPad Prism.

*Heatmaps and Clustering*: Data were analyzed from the the GEO database, accession number GSE185491. rlog normalized counts of RNA sequencing data for NT, RT, and HF samples were produced using the DESeq2 package on the Galaxy web platform public server.^63,64^ Genes of interest from normalized count files were filtered and plotted using the heatmap2 function on Galaxy^64^ as well as the pheatmap R package (version 1.0.12, https://CRAN.R-project.org/package=pheatmap), employing bidirectional complete Euclidian clustering.

*Gene Ontology Analyses*: Data were analyzed from the the GEO database, accession number GSE185491. For both of the pairwise comparison sets of HF vs. NT and RT vs. NT, a list of significantly differentially expressed genes (FDR<0.05) and their respective log_2_(Fold Change) values were uploaded to the PANTHER knowledgebase for gene ontology (GO) analysis.^65^ The PANTHER statistical enrichment test was used to generate lists of significantly upregulated and downregulated GO terms from the biological process, cellular component, and molecular function databases using a threshold of FDR<0.05.^66^ Full data output from GO analyses can be found in **Supplementary Data 1** and **2**.

*YAP1 Interactome*: The YAP1 interactome was generated using the GeneMANIA prediction server plugin in Cytoscape.^52,53^ Differentially expressed nodes from the interactome were highlighted and plotted on a separate heatmap as described earlier in the Heatmaps and Clustering section.

*Statistics*: Statistical analysis was performed in PRISM where t-test (two-tailed) was used to compare two experimental groups and one-way or two-way ANOVA was used for comparison of more than two experimental groups, with p<0.05 considered as significant. Multiple comparisons were performed using Tukey’s test. Normal distribution and homogeneity of variance were tested. When normality was not confirmed, Kruskal-Wallis one-way analysis of variance with Dunn’s multiple comparisons test was performed instead.

## Acknowledgements

We thank K. Chan and J. Moffat for transducing and establishing a stable fluorescent immortalized human podocyte cell line used in the study, C. Ng for fluorescently labelling mouse conditionally immortalized podocyte cell line, R. John for providing insights on histology, L. Fiddes and Y. Chen for sample preparation for SEM imaging, J. Tam and M. Li for technical help with SEM image acquisition, L. Lukic and K. Soon for technical help with microfabrication of thermoplastic substrates. This study is funded by the Natural Sciences and Engineering Research Council of Canada (NSERC) Idea to Innovation grant (I2IPJ 567631-22), NSERC Discovery Grant (RGPIN 326982-10), Canadian Institutes of Health Research (CIHR) Foundation Grant FDN-167274, Center for Research and Applications in Fluidic Technologies Project Award (CRAFT, 512814) and Canada Foundation for Innovation Grant #36442. MR was supported by Killam Fellowship and Canada Research Chair.

## Author contributions

C.L. designed and performed experiments, analyzed data, and prepared the manuscript; P.A., K.T.W., and M.R. analyzed RNA sequencing data; K.T.W. performed EV analysis and western blotting, and contributed to qPCR; S.S.L. performed multifractal and Fourier transform analysis, directionality analysis, and LDH assay, and contributed to fluorescence image analysis; N.R. contributed to qPCR; Y.Z. contributed to RNA sequencing data analysis and fluorescence image acquisition and analysis; K.M. fabricated fractal topographical 96- and 384-well plates; D.B., S.P-G., and S.O. contributed to fluorescence image acquisition; T.C., X.S., L.S., S.R-R., E.V., C.Y.L., A.R., and S.C-F. contributed to review and editing of the manuscript; T.V., M.S., T.F., U.B., and A.K. contributed to conceptualization and co-supervision of the study and review and editing of the manuscript; M.R. conceived and supervised the study, designed experiments, and wrote the manuscript.

## Competing interest

M.R., A.K., and C.L. are inventors on a US Patent application relating to fractal cues for cell maturation.

## Data Availability

The authors declare that the main data supporting the findings of this study are available within the paper and its supplementary information files. Additional data are available from the corresponding author upon reasonable request.

## Supplementary Information

**Supplementary Figure 1.**
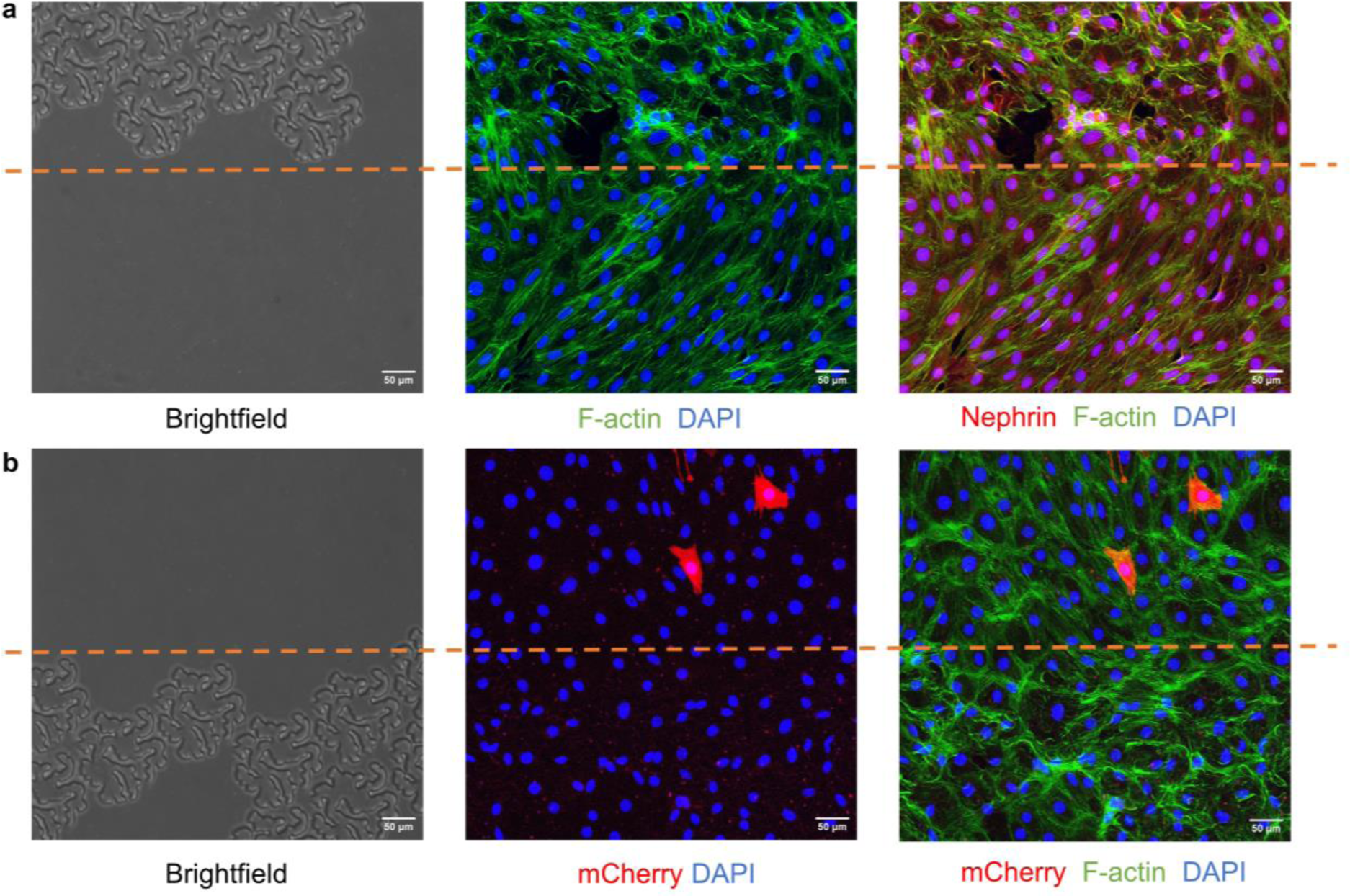
Biomimetic fractal topography guided cytoskeleton arrangement of mouse podocytes. Partially covered fractal patterns under optical microscopy and maximum intensity projection in z-direction of fluorescent stain of f-actin (green), DAPI (blue), and **a,** podocyte specific marker nephrin (red), **b,** cells selectively labelled with mCherry (red), from mouse podocytes cultivated on thermoplastic elastomer substrates partially covered with fractal patterns. DAPI, 4′,6-diamidino-2-phenylindole.

**Supplementary Figure 2.**
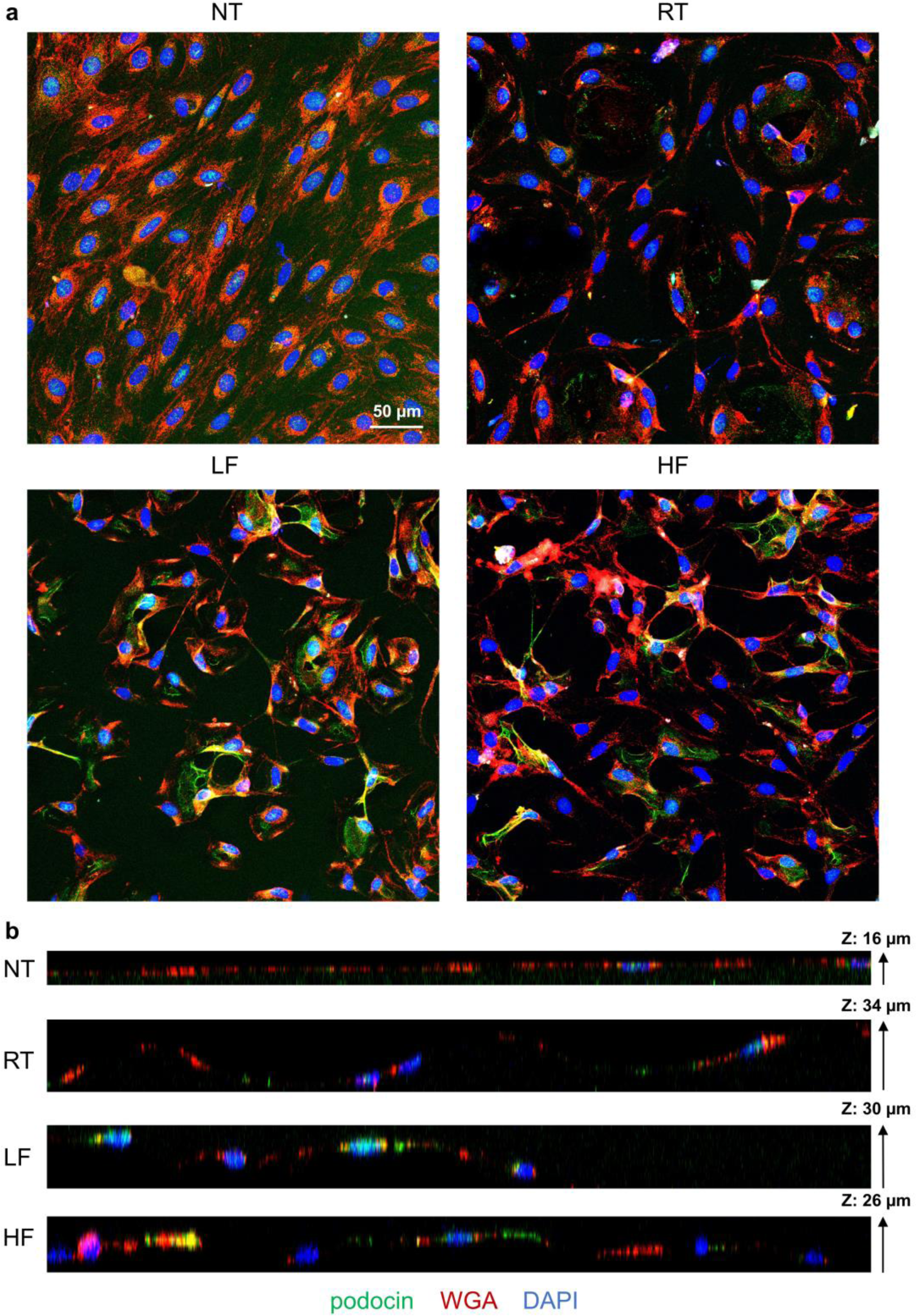
Podocin expression was localized to topographic regions in mouse podocytes cultivated on NT, RT, LF, and HF scaffolds. **a,** Maximum intensity projection in z-direction, and **b,** orthogonal projection of immunofluorescent stain of podocyte specific marker podocin, from mouse podocytes cultivated on flat (NT), round (RT), low-fractal (LF), and high-fractal (HF) topographic substrates. WGA, wheat germ agglutinin; DAPI, 4′,6-diamidino-2-phenylindole.

**Supplementary Figure 3.**
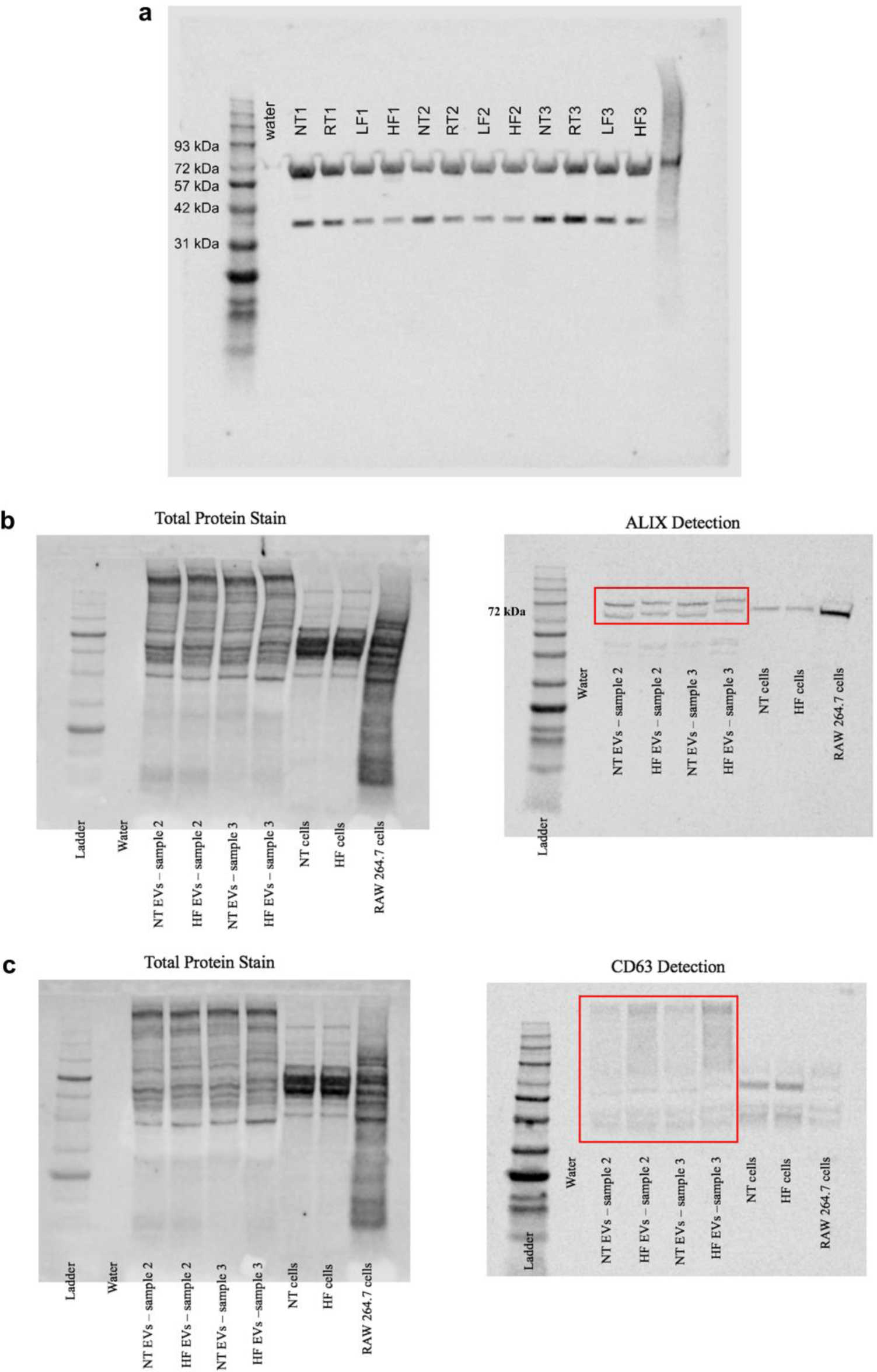
Western blot of podocalyxin and EV markers ALIX and CD63 from mouse podocytes grown on flat and topographical substrates. **a,** Full blot of PODXL (top band) and GAPDH (bottom band) from podocytes cultivated on NT, RT, LF and HF scaffolds. **b-c,** Full blot of total protein stain and **b,** ALIX and **c,** CD63 from EV samples isolated from culture media of mouse podocytes cultivated on NT and HF scaffolds.

**Supplementary Figure 4.**
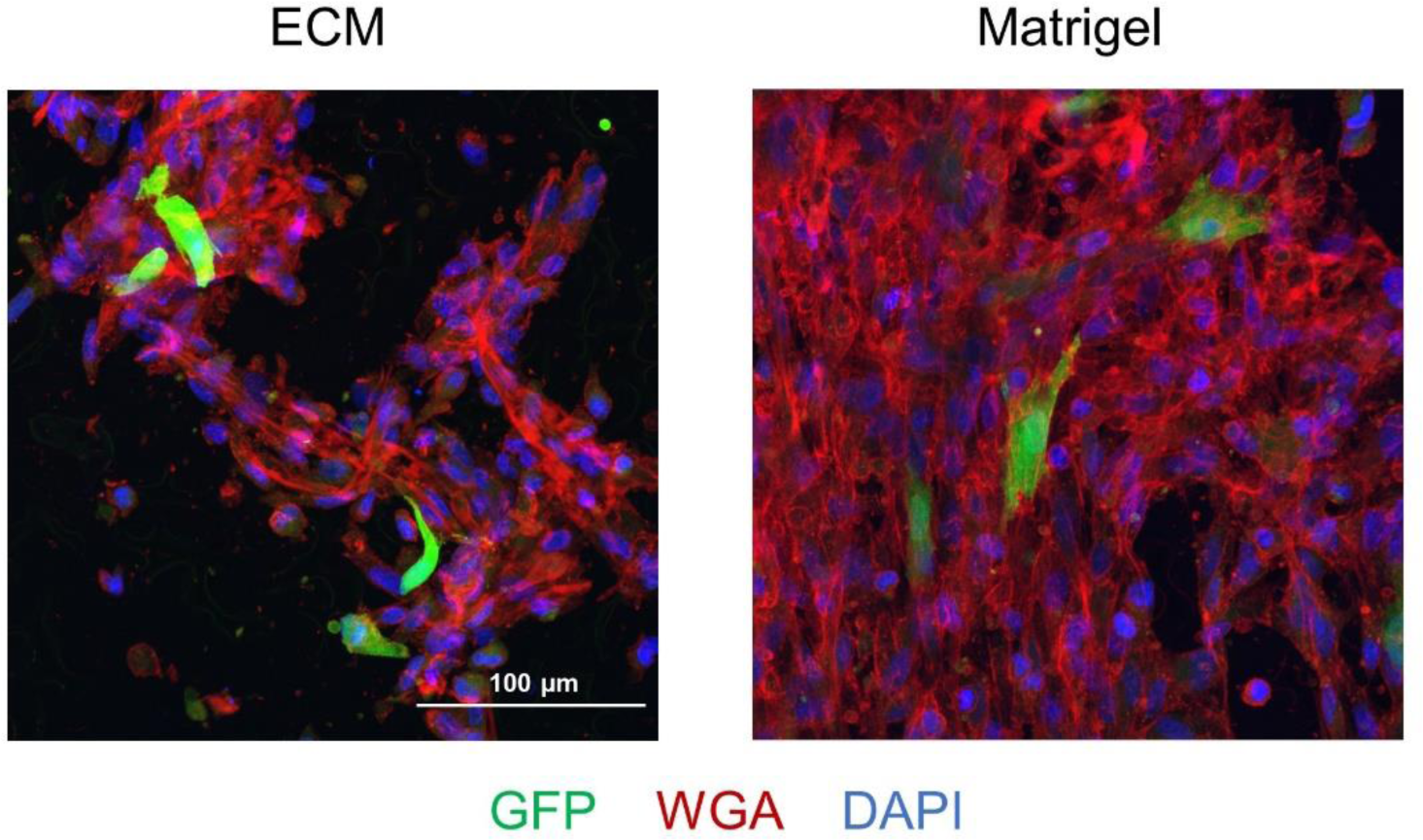
Difference in cell coverage of Mosaic assay human podocyte cell line after 5 days in culture on HF topographical substrates with different coatings. Mosaic assay cells on HF scaffolds coated with NativeCoat ECM vs. Matrigel. HF, high-fractal; GFP, green fluorescent protein; WGA, wheat germ agglutinin; DAPI, 4′,6-diamidino-2-phenylindole.

**Supplementary Figure 5.**
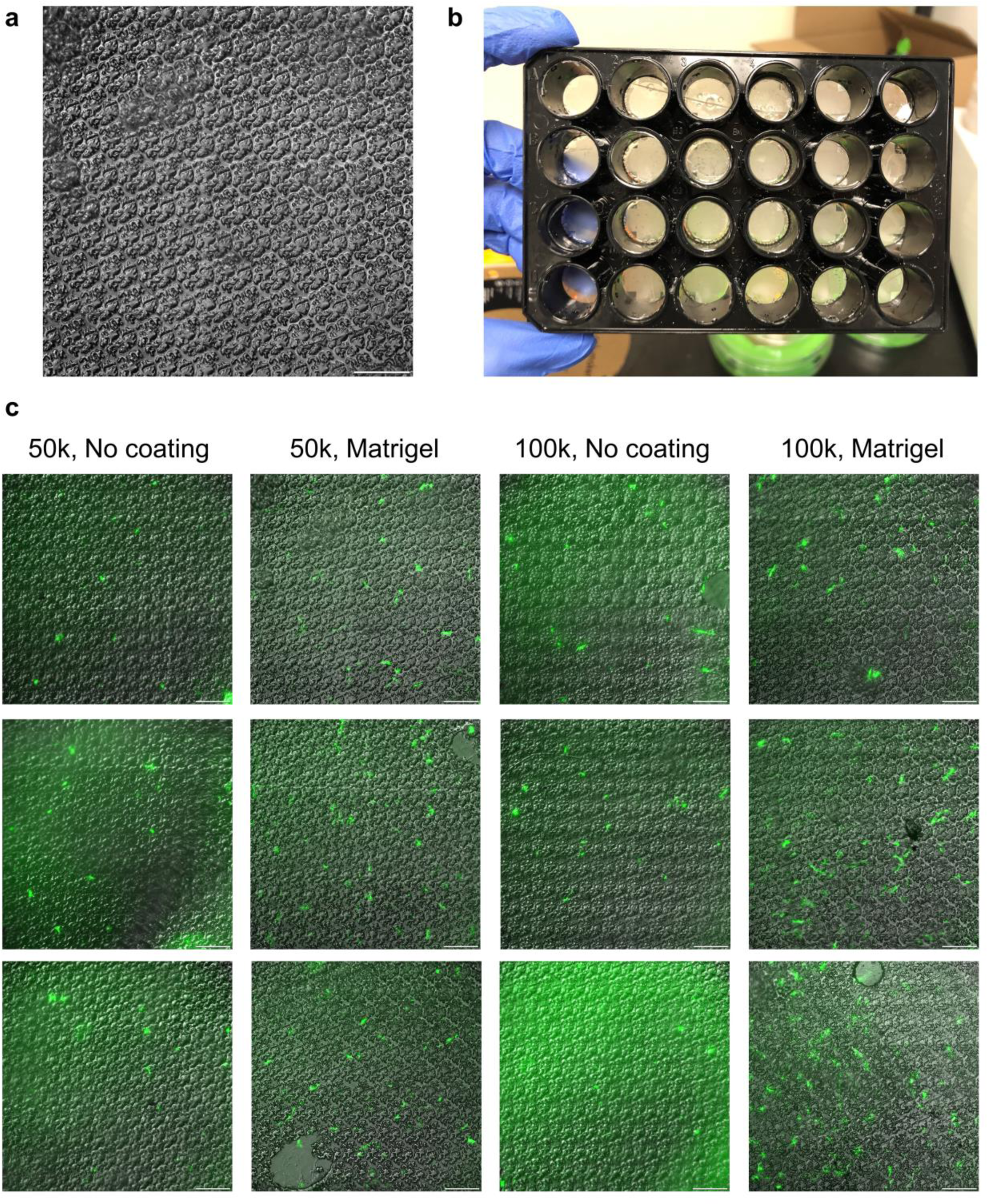
Polystyrene substrates with high density HF patterns. **a,** Hot embossed tissue culture polystyrene substrates with high density HF patterns. **b,** High density HF patterns covering the entire bottom in a 24-well format. **c,** Mosaic assay of non-conditionally immortalized human podocytes after 4 days of cultivation on patterned polystyrene substrates at a seeding density of 50,000 or 100,000 cells/cm^2^ with and without Matrigel coating. Scale bar, 200 µm.

**Supplementary Figure 6.**
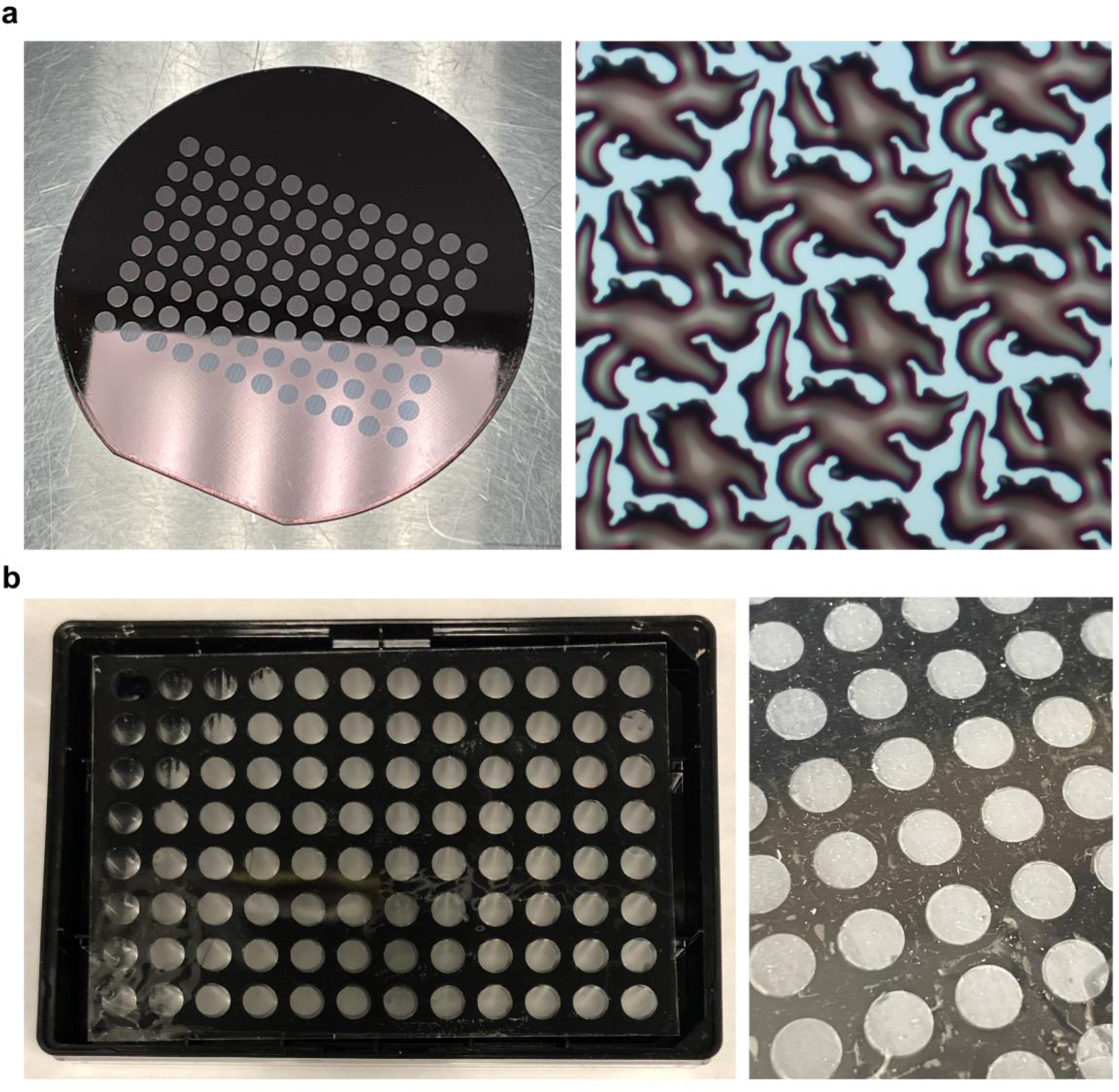
a, Master mold with high density HF pattern in individual well areas. Features were smoothed after reflow. **b,** High density HF patterns covering the entire bottom in a 96-well format.

**Supplementary Figure 7.**
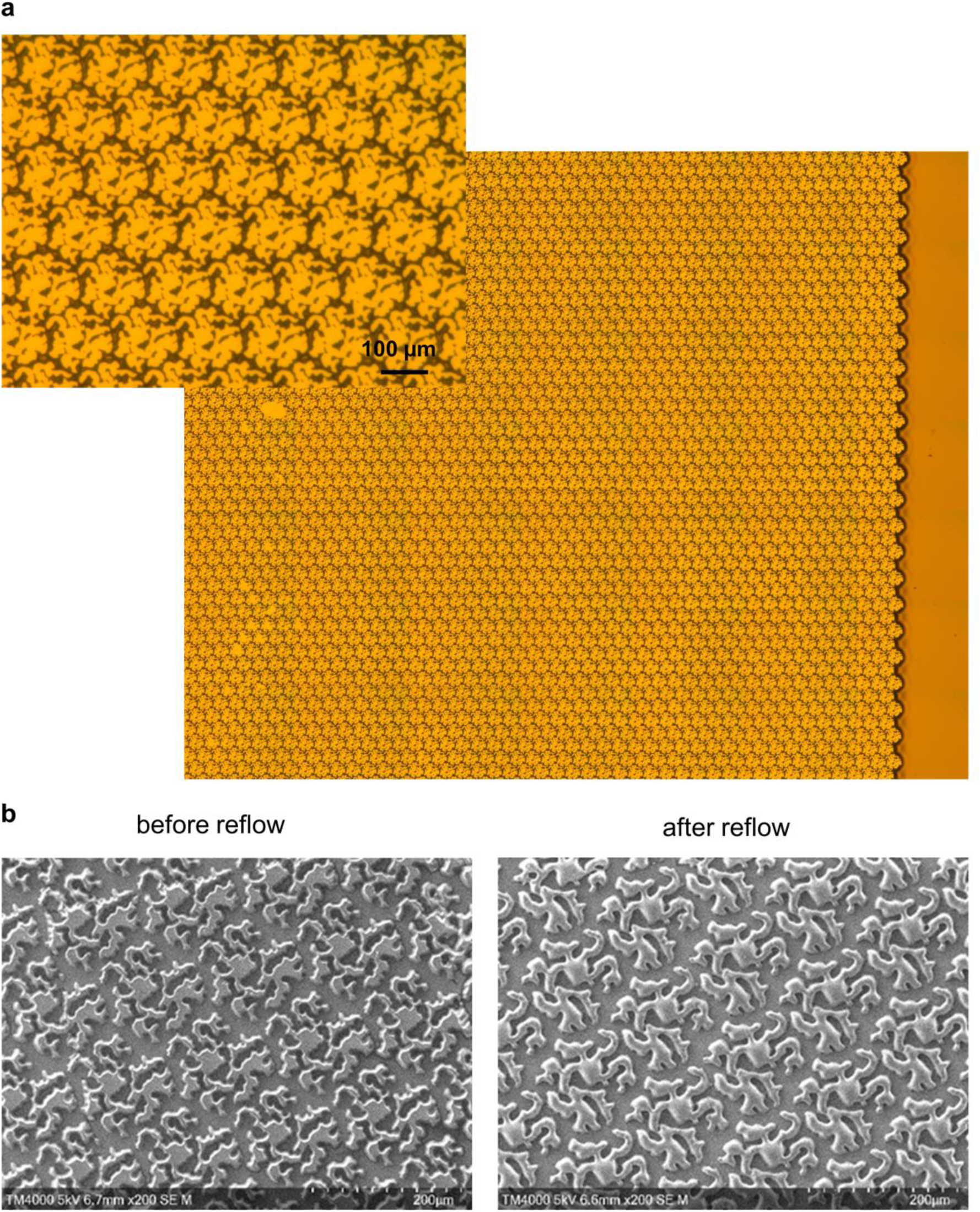
Development of AZ P4620 positive photoresist master mold with densely packed high fractal patterns (high density HF) used in the fabrication of topographic substrates for scalable production of well plate bases with high density HF patterning. **a,** Images of master mold under microscopy at high and low magnifications. **b,** SEM images of a master mold before and after reflow.

**Supplementary Figure 8.**
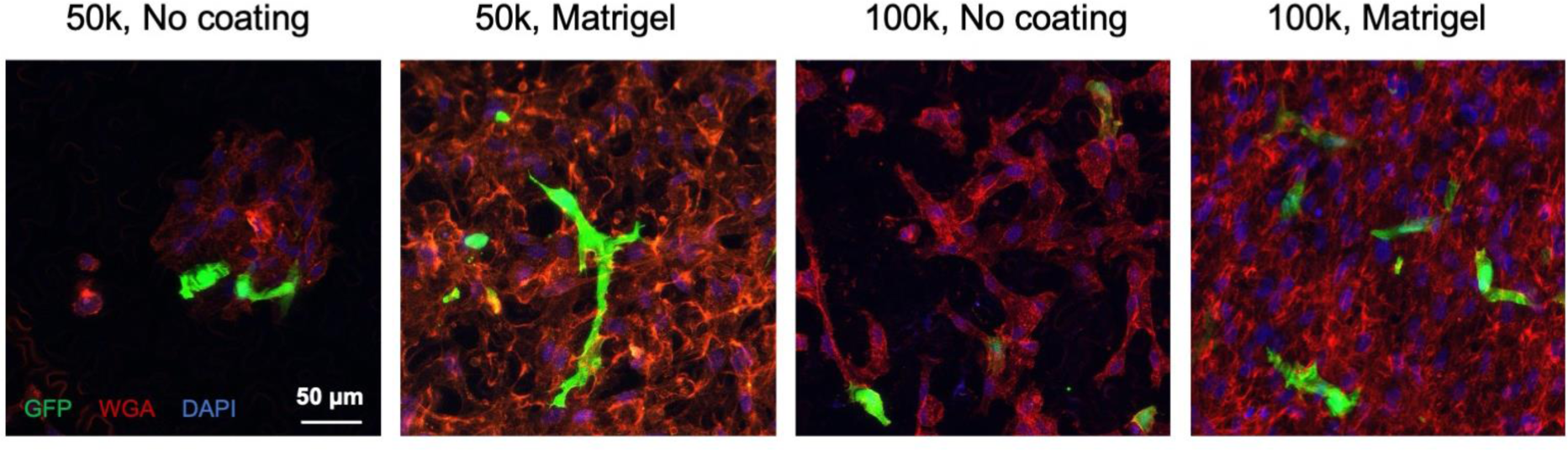
Mosaic human podocytes cultivated for 5 days on TPE fractal topographical 384-well plate with and without Matrigel coating at different seeding densities (50,000 and 100,000 cells/cm^2^). TPE, thermoplastic elastomer.

**Supplementary Figure 9.**
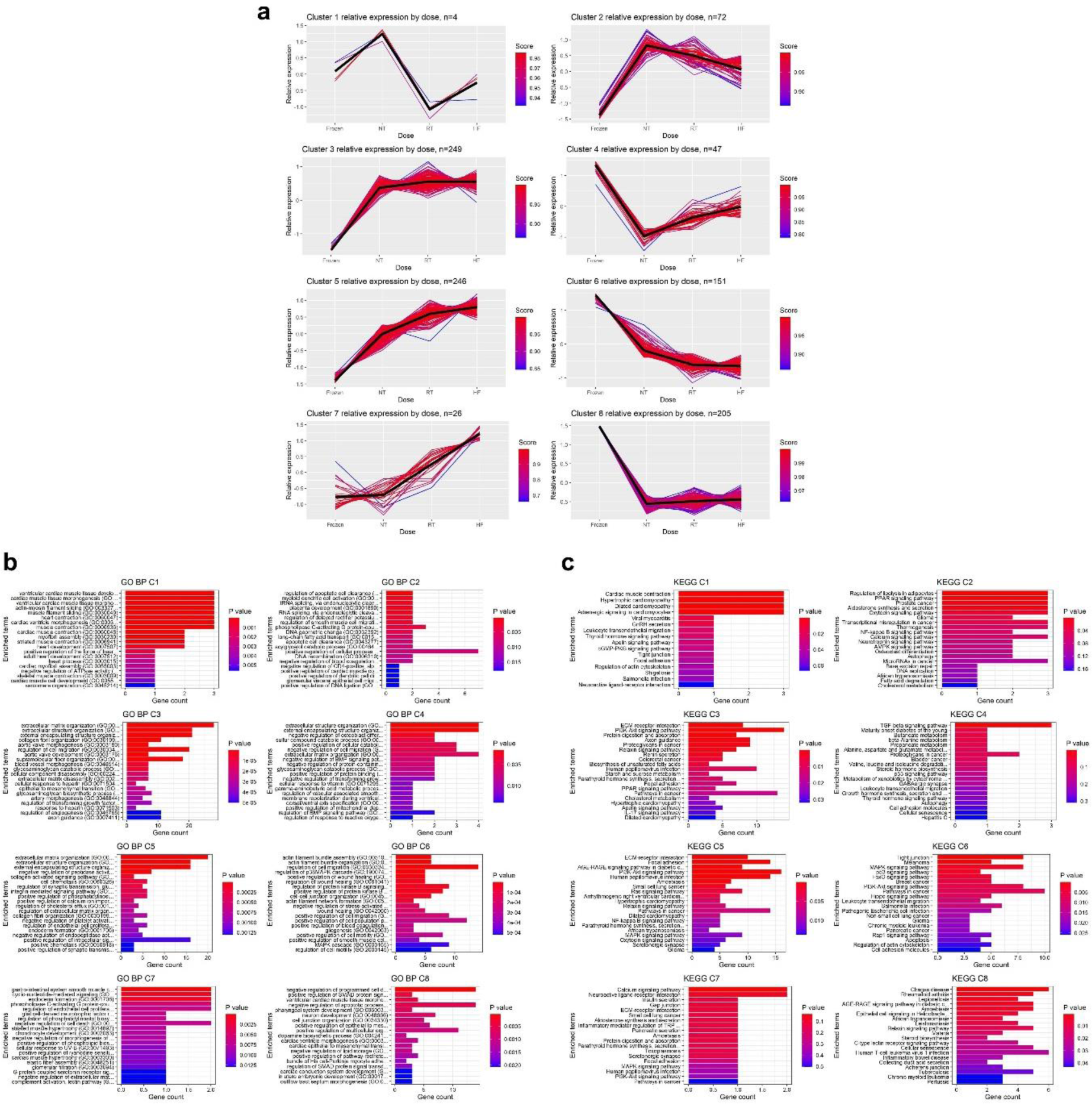
Gene clustering in baseline cells, NT, RT and HF samples. **a,** K-means sub-cluster analysis using 1000 most variable genes. Enrichment analysis on sub-clusters for **b,** Gene ontologies (GO) terms of biological processes and **c,** KEGG analysis by cluster. RNA sequencing data from primary GW18 fetal human podocytes after 3 days in culture on n=4 scaffold samples per group and baseline cells prior to culture available on GEO repository under accession code GSE185491 was used for analysis.^32^ NT, no topography; RT, round topography; HF, high fractal.

**Supplementary Figure 10.**
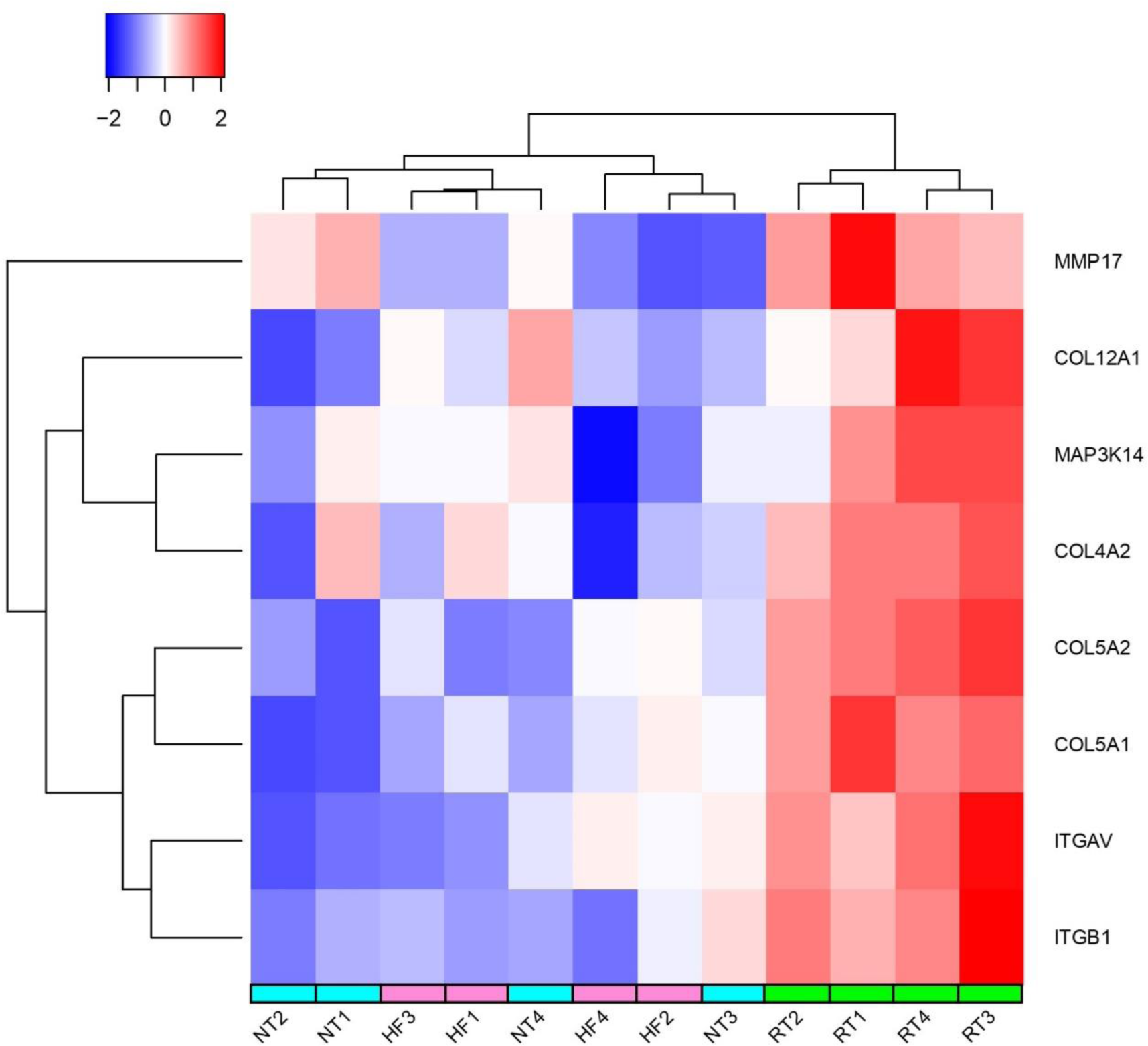
Heatmap of relevant ECM and cell adhesion genes that were significantly different between RT and HF, but not between HF and NT. RNA sequencing data from primary GW18 fetal human podocytes after 3 days in culture on n=4 scaffold samples per group and baseline cells prior to culture available on GEO repository under accession code GSE185491 was used for analysis.^32^ NT, no topography; RT, round topography; HF, high fractal.

**Supplementary Table 1.** List of antibodies and fluorescent stains used in the study.

| Antibody/Stain | Supplier | Dilution |
| --- | --- | --- |
| Podocin polyclonal antibody (host: rabbit) | Abcam, ab50339 | 1:200 |
| Nephrin polyclonal antibody (host: rabbit) | Invitrogen, PA5-20330 | 1:500 |
| Nestin monoclonal antibody (host: mouse) | Invitrogen, MA1-110 | 1:500 |
| Alexa Fluor 488 Phalloidin | Invitrogen, A12379 | 1:500-1:1000 |
| Rhodamine labeled Wheat Germ Agglutinin | Vector Laboratories, RL-1022 | 1:500 |
| Goat anti-rabbit IgG FITC | LifeTech A16105 | 1:500 |
| Goat anti-rabbit IgG Alexa Fluor 647 | Invitrogen A21245 | 1:500 |
| Goat anti-mouse IgG Alexa Fluor 568 | Invitrogen A11004 | 1:500 |

**Supplementary Table 2.**
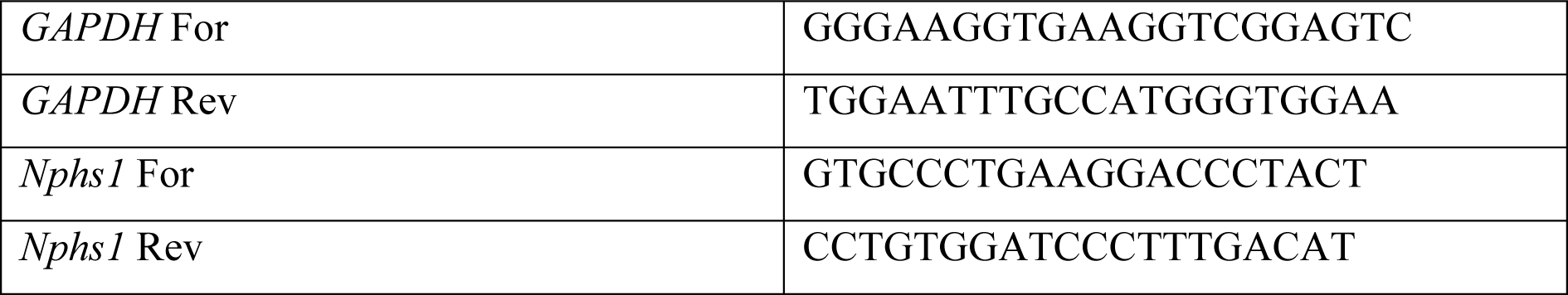
List of primers used for qPCR.

## Notes

https://www.ncbi.nlm.nih.gov/geo/query/acc.cgi?acc=GSE185491

